# Altered GABAergic inhibition in CA1 pyramidal neurons modifies despair-like behavior in mice

**DOI:** 10.1101/2020.02.18.953786

**Authors:** Sang Ho Yoon, Geehoon Chung, Woo Seok Song, Sung Pyo Oh, Jeongjin Kim, Sang Jeong Kim, Myoung-Hwan Kim

## Abstract

Despair is a common human feeling characterized by the loss of hope and is a core symptom of depressive disorders. However, little is known regarding neural circuits mediating despair and their modulation by antidepressants. Here we show that alterations in inhibitory synaptic transmission in the hippocampus affect behavioral despair in mice. Reduced interneuron density, knockdown of GABA_A_ receptor gamma 2 subunit gene (*Gabrg2*) or DREADD-mediated suppression of interneuron activity resulted in disinhibition of CA1 neurons and anti-despair-like behaviors in mice. Similarly, a low dose of pentylenetetrazol, a GABA_A_R antagonist, induced transient anti-despair-like behaviors, with rapid eukaryotic elongation factor 2 (eEF2) activation in the hippocampus. Conversely, pharmacological and chemogenetic potentiation of GABAergic transmission in CA1 neurons induced despair-like behaviors. The antidepressant ketamine rapidly increased c-Fos expression in CA1 neurons and induced anti-despair-like behaviors. These results suggest that the enhanced hippocampal CA1 neuron activity induces anti-despair-like behaviors and contributes to the antidepressant effects of ketamine.

## Introduction

Despair is a common human feeling characterized by the loss of hope that significantly disrupts motivation and goal-directed thinking^1^. Notably, despair is one of the core symptoms of depressive disorders, and hopelessness is a strong risk factor for suicidal thoughts^2, 3^. Although the brain mechanisms underlying despair remain unknown, stress is a potent risk factor for depressive symptoms and behavioral despair in humans and laboratory animals, respectively^3, 4^. Conversely, antidepressants improve despair and suicidal thoughts in patients with depression, as well as behavioral despair in rodent models^3–5^.

A growing body of evidence suggests that rapid-acting antidepressant agents, such as ketamine (an NMDA receptor blocker), the ketamine metabolite (2R,6R)-hydroxynorketamine, and scopolamine (a muscarinic acetylcholine receptor antagonist), share core mechanisms for synaptic action by rapidly suppressing synaptic inhibition to pyramidal neurons in the medial prefrontal cortex (mPFC) and hippocampus^6, 7^. Enhanced glutamatergic neurotransmission, termed the “glutamate burst” and caused by disinhibition of pyramidal neurons, is thought to trigger protein synthesis-dependent LTP (long-term potentiation)-like changes in synaptic efficacy and neuronal morphology in the brain areas implicated in depressive disorders^4^. The activation of signaling cascades associated with protein synthesis, including eukaryotic elongation factor 2 (eEF2), mammalian target of rapamycin (mTOR), and extracellular signal-regulated kinase (ERK), reportedly increases brain-derived neurotrophic factor (BDNF) and synaptic protein synthesis, modifies the function of neural circuits, and mediates the effects of rapid-acting antidepressant agents^4, 8, 9^.

A low dose of ketamine rapidly disinhibits hippocampal CA1 neurons, consequently increasing the probability of synaptically driven action potential in CA1 pyramidal neurons^7^. In line with these findings, L-655,708 or MRK-016, negative allosteric modulators of the α5-subunit-containing GABA_A_ receptor (GABA_A_R), rapidly reverse depression-like behaviors and stress-impaired AMPAR-mediated transmission at hippocampal temporoammonic-CA1 synapses in stressed rodents^10^. Collectively, these findings demonstrate that disinhibition of CA1 pyramidal neurons might be associated with the effects of rapid-acting antidepressants.

We have previously reported that deficiency of RalBP1, a downstream effector of the small GTP-binding proteins RalA and RalB, results in reduced interneuron density in the hippocampal CA1 areas^11^. Basal excitatory synaptic transmission and LTP at the Schaffer collateral-CA1 synapse were normal in RalBP1^-/-^ mice, but long-term depression (LTD) was modestly attenuated^12^. Based on these findings, we investigated whether reduced synaptic inhibition in CA1 pyramidal neurons caused by reduced interneuron numbers in RalBP1-mutant mice recapitulates disinhibition and behavioral effects of rapid-acting antidepressants.

Although depression- or stress-related synaptic deficits and antidepressant-induced synaptic changes are observed in both the mPFC and the hippocampus, the behavioural responses to altered activity of hippocampal CA1 microcircuits remain unclear. To address this issue, we selectively manipulated the activity of CA1 interneurons or inhibitory synaptic transmission to CA1 pyramidal neurons. We show here that disinhibition of CA1 pyramidal neurons is sufficient to induce anti-despair-like behaviors in mice. The present study suggests that the enhanced hippocampal CA1 neuron activity contributes to the antidepressant effects of ketamine.

## Results

### Reduced synaptic inhibition in CA1 neurons and anti-despair-like behaviors in RalBP1^-/-^ mice

RalBP1 is widely expressed in brain neurons and RalBP1^-/-^ mice exhibit fewer GAD67 (glutamic acid decarboxylase 67)-positive neurons in the hippocampus^11, 12^. To examine the functional consequence of reduced interneurons in the hippocampal CA1 area, we measured the ratio of synaptic excitation (E) and inhibition (I) to CA1 pyramidal neurons. We first recorded evoked excitatory postsynaptic currents (EPSCs) at the reversal potential of inhibitory postsynaptic currents (IPSCs) using a low-chloride (15 mM) pipette solution and then recorded IPSCs at the reversal potential of EPSCs in the same cell (Fig. 1a). The reversal potentials of IPSCs and EPSCs were −57 mV and +3 mV, respectively (Supplementary Fig. 1). The amplitudes of evoked IPSCs were significantly decreased in RalBP1^-/-^ CA1 neurons, whereas evoked EPSCs were unaltered (Fig. 1b). These results indicate that the RalBP1^-/-^ mutation leads to an increased synaptic E/I ratio in CA1 neurons (Fig. 1c).

**Fig. 1:**
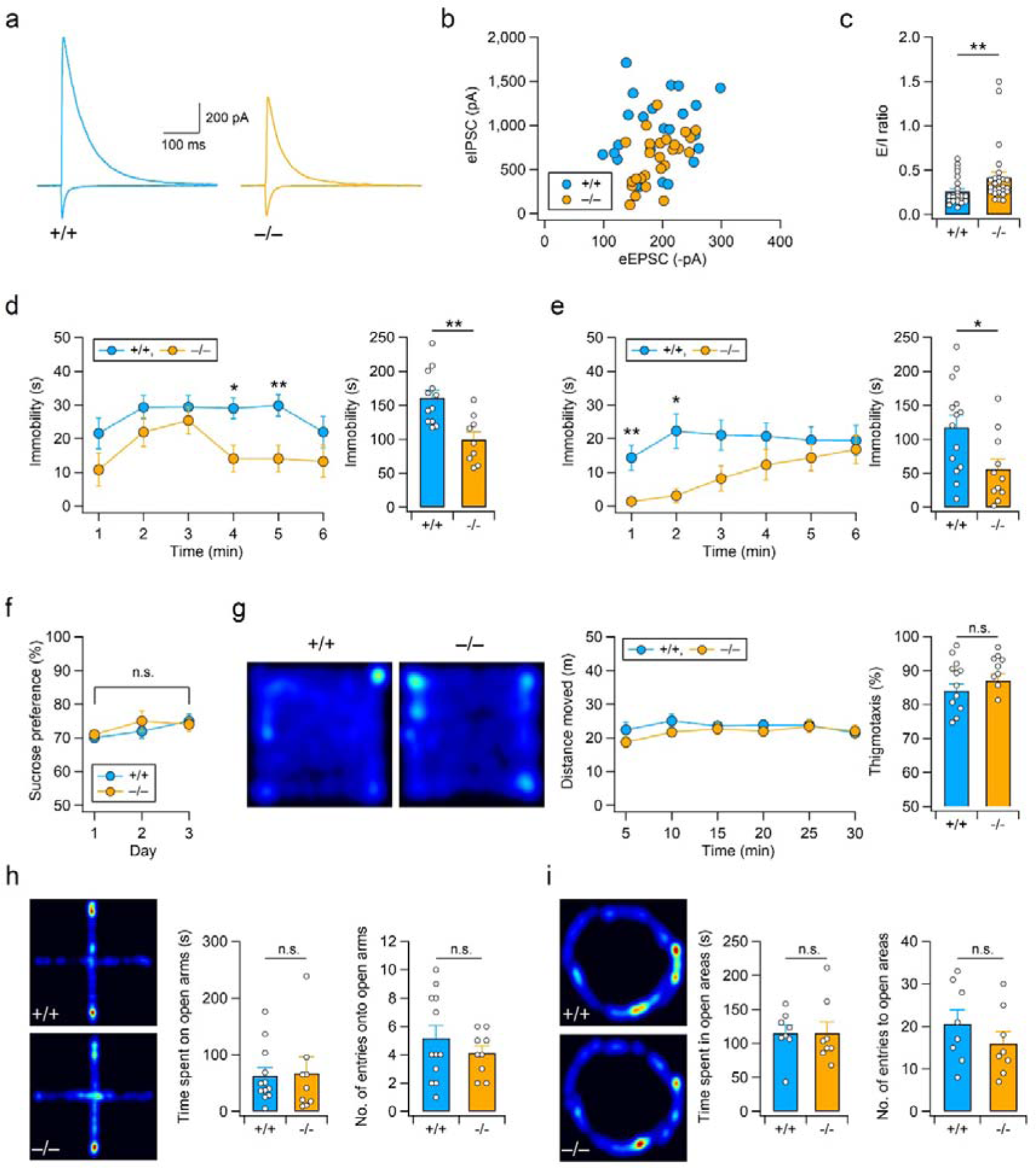
RalBP1^-/-^ mice display decreased synaptic inhibition and anti-despair-like behaviors but normal anxiety levels. **(a-c)** Enhanced excitation-inhibition (E/I) ratio in RalBP1^-/-^ CA1 pyramidal neurons. (a) Representative traces (average of 15 consecutive sweeps) of evoked EPSCs (downward deflections at −57 mV) and IPSCs (upward deflections at +3 mV) recorded in CA1 pyramidal neurons from WT (RalBP1^+/+^) and RalBP1^-/-^ mice. (b) The mean amplitude of evoked IPSCs measured in each cell was plotted against the mean amplitude of EPSCs. (c) The E/I ratios are significantly increased in RalBP1^-/-^ mice. (d, e) RalBP1^-/-^ mice display anti-despair-like behavior. (d) Immobility time of mice was measured every 1 min during the TST (left). Total time spent immobile over 6 min in the TST (right). (e) Forced swimming test (FST) of RalBP1^+/+^ and RalBP1^−/−^ mice (left). RalBP1^−/−^ mice spent significantly less time immobile during the 6-min FST (right). (f) Sucrose preference during the 3-day-sucrose-consumption test is unaltered in RalBP1^−/−^ mice. (g-i) Normal anxiety levels and locomotor activity in RalBP1^−/−^ mice. (g) Representative activity paths (left), distance moved (middle), and the thigmotaxis (a tendency to remain close to walls) levels (right) in the open-field test. (h) Representative activity traces (left), time spent in the open arms (middle), and the number of open-arm entries (right) of WT and RalBP1^−/−^ mice in the elevated plus maze test. (i) Representative activity traces (left), time spent in the open areas (middle), and the number of entries into open areas (right) of WT and RalBP1^−/−^ mice in the elevated zero-maze test.

The hippocampus contains a mixed population of GABAergic (γ-aminobutyric acid-ergic) interneurons, derived from the medial ganglionic eminence (MGE) and caudal ganglionic eminence (CGE)^13^. We observed fewer MGE-derived parvalbumin (PV)-expressing interneurons, but a normal density of somatostatin (SST)-expressing interneurons, which originate from both MGE and CGE, in the RalBP1^-/-^ hippocampal CA1 region (Supplementary Fig. 2a, b). On examining the density of CGE-derived interneurons expressing vasoactive intestinal peptide (VIP) and/or cholecystokinin (CCK), the density of VIP-positive interneurons, but not CCK-positive interneurons, was reduced in the RalBP1^-/-^ CA1 area (Supplementary Fig. 2c, d).

To understand the behavioral consequences of reduced synaptic inhibition in RalBP1^-/-^ mice, we first investigated mood-related behaviors. We employed the tail suspension test (TST) and forced swim test (FST) to examine depression-like behaviors in RalBP1^-/-^ mice^14, 15^. Compared with the wild-type (WT) control, RalBP1^-/-^ mice spent significantly less time immobile during the TST and FST (Fig. 1d,e), suggesting that RalBP1 deficiency reduces behavioral despair in mice. We examined additional depression-related behaviors, including anhedonia, locomotor activity, and anxiety, using a series of behavioral tests. During the sucrose consumption test, the degree of preference for sucrose was indistinguishable between genotypes (Fig. 1f). RalBP1^-/-^ mice exhibited normal behavior in an open field box (Fig. 1g). Furthermore, RalBP1^-/-^ mice did not present behavioral signs of seizures or epileptic electroencephalogram (EEG) rhythms in their homecages (Supplementary Fig. 3). These observations exclude the possibility that the reduced immobility of RalBP1^-/-^ mice during both TST and FST might stem from behavioral hyperactivity^16^ or epilepsy. Consistent with normal levels of thigmotaxis in the open-field test (Fig. 1g), RalBP1^-/-^ mice exhibited normal anxiety levels in both the elevated plus-maze and zero-maze tests (Fig. 1h, i). Collectively, these results suggest that the deficiency of RalBP1 selectively affects despair-like behaviors in mice.

### RalBP1 deficiency does not affect hippocampus-dependent learning, social interaction and sensory gating

We examined hippocampus-dependent learning behaviors in RalBP1^-/-^ mice. During the Morris water maze test, both genotypes showed similar escape latencies and quadrant occupancy (Supplementary Fig. 4a-c). Performance in the Morris water maze was not affected by motor functions in RalBP1^-/-^ mice, as revealed by normal swimming speed (+/+, 16.54 ± 1.29 cm/s; –/–, 17.15 ± 1.37 cm/s; P > 0.05 by Student’s t-test) and rotarod performance (Supplementary Fig. 4d). RalBP1^-/-^ mice also performed normally in both the contextual and cued fear conditioning tests (Supplementary Fig. 4e). These results suggest that hippocampus-dependent learning and memory remain normal in RalBP1^-/-^ mice.

As dysfunction of GABAergic neurotransmission has been implicated in the pathophysiology of autism and schizophrenia^17^, we assessed social interaction, repetitive behaviors, and sensory gating function in RalBP1^-/-^ mice. In the three-chamber test^18^, WT and RalBP1^-/-^ mice showed similar sociability and preference for social novelty (Supplementary Fig. 4f-i). In addition, RalBP1^-/-^ mice did not display repetitive or stereotyped behaviors (Supplementary Fig. 4j), as determined by normal behaviors in the marble-burying test^19^. Impairment in prepulse inhibition of the acoustic startle response, a deficit in sensorimotor gating, which is frequently observed in animal models of schizophrenia^20^, was not detected in RalBP1^-/-^ mice (Supplementary Fig. 4k).

### Altered hippocampal circuit function is associated with anti-despair-like behavior in RalBP1^-/-^ mice

We wondered whether changes in brain regions other than the hippocampus are responsible for anti-despair-like behaviors in RalBP1^-/-^ mice. The PFC and amygdala have been implicated in despair/depression-like behaviors^21–24^. However, the interneuron density in the mPFC and basolateral amygdala of RalBP1^-/-^ mice were comparable with those of WT mice (Supplementary Fig. 5).

Based on the observed altered hippocampal network and anti-despair-like behaviors in RalBP1^-/-^ mice, we wondered whether pharmacological potentiation of GABAergic transmission in the hippocampal CA1 area would induce despair-like behavior in mice. Indeed, bilateral infusion of muscimol, a GABA_A_R agonist, into the dorsal hippocampal CA1 region of WT mice induced despair-like behaviors in a dose-dependent manner during the TST (Fig. 2a, b). We further examined whether enhancement of GABAergic transmission in the CA1 area would reverse anti-despair-like behavior in RalBP1^-/-^ mice. Cannula implantation did not affect anti-despair-like behaviors in RalBP1^-/-^ mice, as reduced immobility in RalBP1^-/-^ mice was again observed during the TST (Fig. 2c, TST1). However, on infusing muscimol (0.5 µg) bilaterally into the hippocampal CA1 region, the immobility observed in RalBP1^-/-^ mice was comparable with that of saline-treated WT mice (Fig. 2c, TST2). Behavioral changes in RalBP1^-/-^ mice were treatment-specific as RalBP1^-/-^ mice regained anti-despair-like behaviors when both genotypes were administered saline (Fig. 2c, TST3). These results indicate that altered activity in the hippocampal network is responsible for anti-despair-like behaviors in RalBP1^-/-^ mice.

**Fig. 2:**
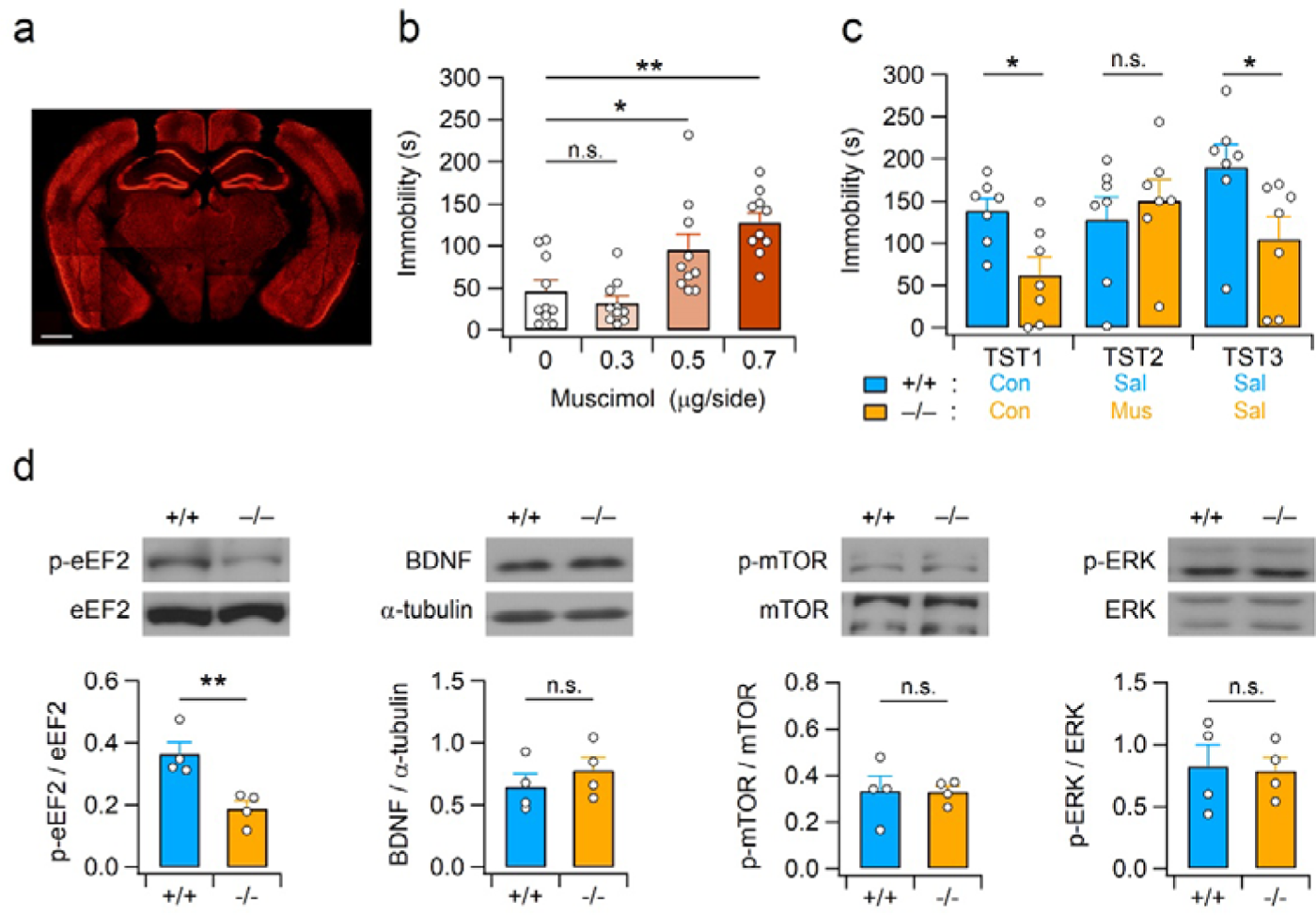
Attenuation of anti-despair-like behavior by intrahippocampal infusion of muscimol and altered hippocampal eEF2 signaling in RalBP1^-/-^ mice. (a) Representative image showing the cannula placement in the mouse brain for muscimol infusion. The section was immunostained with anti-NeuN antibodies. Scale bar, 1 mm. (b) Hippocampal infusion of muscimol increases TST immobility time in a dose-dependent manner. (c) Potentiation of GABAergic transmission in the hippocampal CA1 area reverts the anti-despair-like behavior of RalBP1^-/-^ mice. TST was performed 30 min after completion of infusion and inter-test intervals were 2 days. Con, Sal, and Mus represent no infusion, saline infusion, and muscimol infusion, respectively. (d) Representative western blot images (top) and quantification (bottom) showing enhanced eEF2 activation but normal BDNF, mTOR, and ERK signaling in the RalBP1^-/-^ hippocampus. The blots were cropped, and full blots are presented in Supplementary Fig. 13. Western blotting using anti-α-tubulin antibody was performed to ensure equal protein loading and transfer, and quantification of BDNF levels in each sample.

Activation of signaling pathways associated with protein synthesis, such as the mTOR, ERK and/or eEF2 signaling pathways, is required to mediate behavioral effects of the rapidly acting antidepressant ketamine^8, 9^. We investigated whether activation of these signalling pathways is accompanied in the RalBP1^-/-^ hippocampus. RalBP1^-/-^ mice exhibited significantly reduced levels of phosphorylated (inactivated) eEF2 compared with WT mice, whereas both genotypes showed similar levels of total eEF2 in the hippocampus (Fig. 2d). As dephosphorylation (activation) of eEF2 stimulates protein synthesis and blockade of action potential-mediated network activity promotes phosphorylation of eEF2^9, 25^, enhanced hippocampal network activity caused by disinhibition promotes dephosphorylation of eEF2 and translation of downstream signaling molecules, including BDNF^9^. However, hippocampal BDNF levels in RalBP1^-/-^ mice did not differ from those in WT littermates. Consistent with this result, modification of mTOR and ERK signaling, downstream pathways of BDNF-TrkB (tropomyosin receptor kinase B) signaling, was not detected in the RalBP1^-/-^ mouse hippocampus, as neither total protein expression nor phosphorylation levels differed between genotypes (Fig. 2d). Considering that excitatory synaptic functions, including NMDAR- and AMPAR-mediated synaptic transmissions, as well as expression levels of excitatory synaptic proteins, are normal in the RalBP1^-/-^ mouse hippocampus^12^, activation of the hippocampal eEF2 signaling pathway might be associated with reduced synaptic inhibition in RalBP1^-/-^ mice.

### Suppression of synaptic inhibition in CA1 pyramidal neurons induces anti-despair-like behavior

Synaptic GABA_A_Rs contain one of the three γ subunits (γ_1_ – γ_3_), and the γ_2_ subunit encoded by the Gabrg2 gene plays a critical role in clustering major postsynaptic GABA_A_Rs^26, 27^. To examine the direct causal relationship between reduced synaptic inhibition in CA1 neurons and anti-despair-like behaviors in mice, we prepared four different knockdown vectors containing short hairpin RNA (shRNA) targeting the GABA_A_R γ_2_ subunit (Gabrg2), and determined the gene knockdown efficiency of each construct in HEK293T cells (Supplementary Fig. 6). An shRNA sequence with the highest target knockdown activity (Gabrg2-shRNA#3, shGabrg2) was subcloned into the lentiviral transfer vector and delivered into CA1 neurons (Fig. 3a, b and Supplementary Fig. 6d). Gabrg2 knockdown significantly decreased the frequency but not the amplitude of miniature IPSCs (mIPSCs) in CA1 neurons (Fig. 3c-e). In contrast to mIPSCs, neither the frequency nor amplitude of miniature EPSCs (mEPSCs) was affected by Gabrg2 knockdown (Fig. 3f-h). These results rule out nonspecific neuronal defects induced by shRNA expression or lentivirus infection, excluding homeostatic downregulation of glutamatergic neurotransmission, which is observed in Gabrg2-mutant mice^28^. Normal mIPSCs and mEPSCs in CA1 neurons expressing a non-targeting shRNA (shNT) further support the specificity of Gabrg2 knockdown. Consistent with fewer mIPSCs, CA1 pyramidal neurons infected with shGabrg2 exhibited reduced amplitudes of evoked IPSCs and increased EPSC/IPSC ratios when compared with those infected with shNT or those non-infected (Fig. 3i-l).

**Fig. 3:**
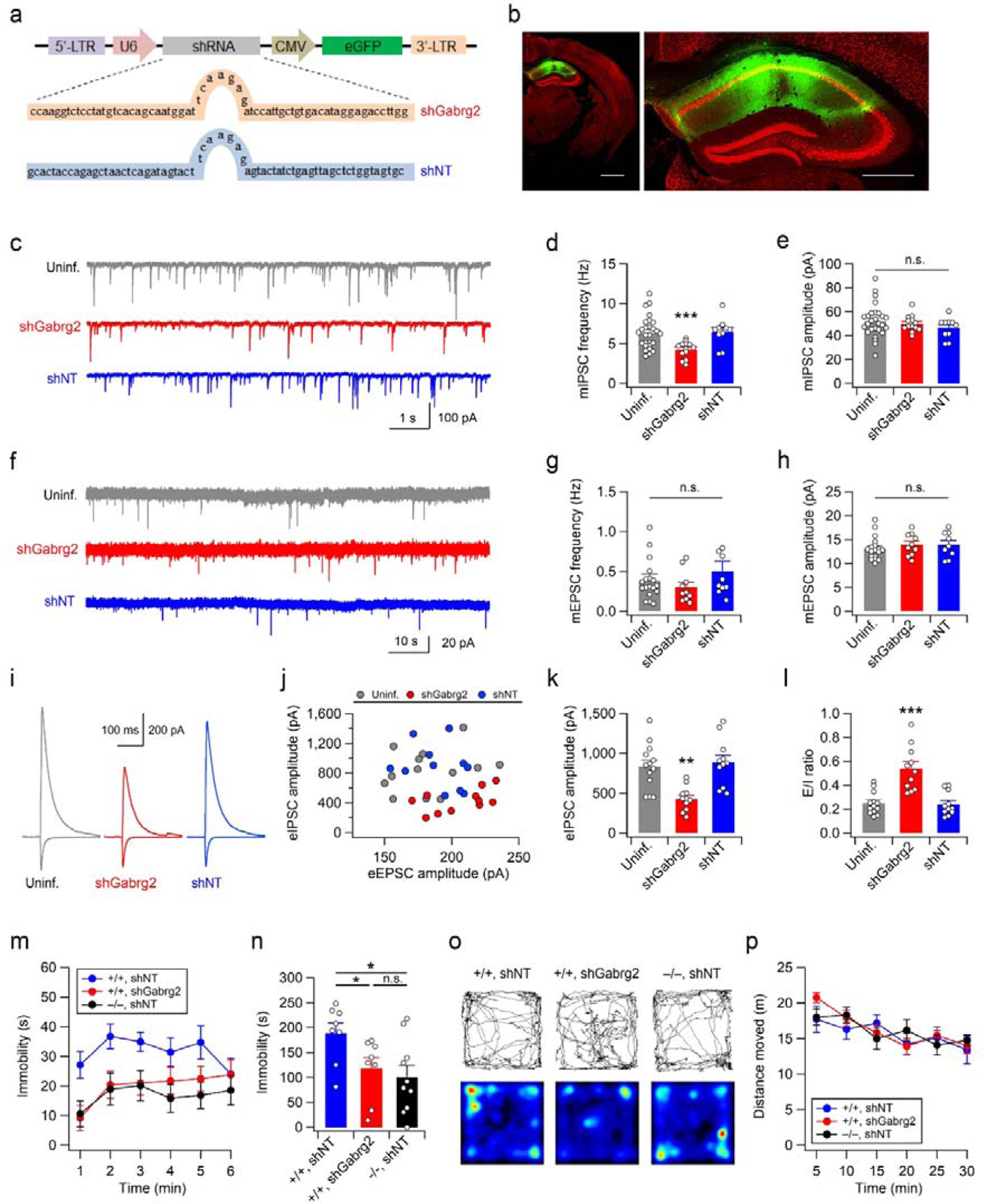
Knockdown of Gabrg2 in hippocampal CA1 neurons reduces despair-like behaviors. (a) Schematic diagram of the viral genome expressing Gabrg2-shRNA#3 (shGabrg2) or non-targeting shRNA (shNT). (b) Coronal brain section from mice expressing eGFP (green) in the hippocampal CA1 area (left). Sections were co-stained with the neuronal marker NeuN (red). Right, enlarged image of the hippocampal area. Scale bars, 1 mm (left) and 500 µm (right). (c) Sample traces of mIPSCs recorded in CA1 pyramidal neurons infected with virus expressing shGabrg2 or NT, and nearby uninfected (uninf.). (d) Gabrg2 knockdown reduces the frequency of mIPSCs. (e) Summary of the mean amplitudes of mIPSCs obtained from uninfected and virus-infected (shGabrg2 or shNT) CA1 neurons. (f-h) Gabrg2 knockdown does not affect mEPSCs. Sample traces (f), mean frequencies (g) and mean amplitudes (h) of mEPSCs recorded in CA1 pyramidal neurons infected with virus expressing shGabrg2 or NT, and nearby uninfected. (i) Representative traces of evoked IPSCs (upward deflections at +3 mV) and EPSCs (downward deflections at −57 mV) recored from CA1 pyramidal neurons. (j) The peak amplitudes of IPSCs are plotted against peak EPSC amplitudes. (k, l) Summary of peak eIPSC amplitudes (k) and the eEPSC-eIPSC (E/I) ratios (l) in the CA1 pyramidal neurons infected with virus expressing shGabrg2 or NT, and nearby uninfected. (m) shRNA-mediated suppression of Gabrg2 expression in hippocampal CA1 neurons reduces immobility of mice in the TST. (n) Summary of time spent immobile in each experimental group during the 6-min TST. (o) Representative images showing exploration path during the first 5 min (top) and entire 30-min period (bottom) in the open field test. (p) Quantification of distance moved in the open field box.

We then examined the effect of Gabrg2 knockdown in CA1 neurons on despair-like behaviors. Similar to RalBP1^-/-^ mice injected with virus expressing shNT, WT mice infected with shGabrg2-expressing virus spent a significantly reduced time immobile in the TST than those infected with shNT virus (Fig. 3m, n). Given that general locomotor activity in the open field box was unaltered by Gabrg2 knockdown in CA1 neurons (Fig. 3o, p), reduced immobility in the TST reflects anti-despair-like behavior in mice.

### Pharmacogenetic manipulation of CA1 interneuron activity modulates behavioural despair

Reduction of CA1 interneurons or inhibitory synaptic transmission to CA1 pyramidal neurons induced anti-despair-like behaviors in mice. Based on these findings, we hypothesized that changes in CA1 interneuron activity might modulate behavioral despair. To selectively manipulate the activity of CA1 interneurons, we expressed excitatory (hM3Dq) or inhibitory (hM4Di) designer receptors exclusively activated by designer drugs (DREADDs) in the hippocampal CA1 area of Vgat-ires-Cre knock-in (Vgat-Cre) mice using adeno-associated virus (AAV) carrying Cre-dependent (double-floxed inverse orientation, DIO) DREADD receptor expression vectors (Fig. 4a). mCherry signals were restricted to GAD67-positive inhibitory neurons (Fig. 4b), including PV- and SST-expressing neurons (Supplementary Fig. 7).

**Fig. 4:**
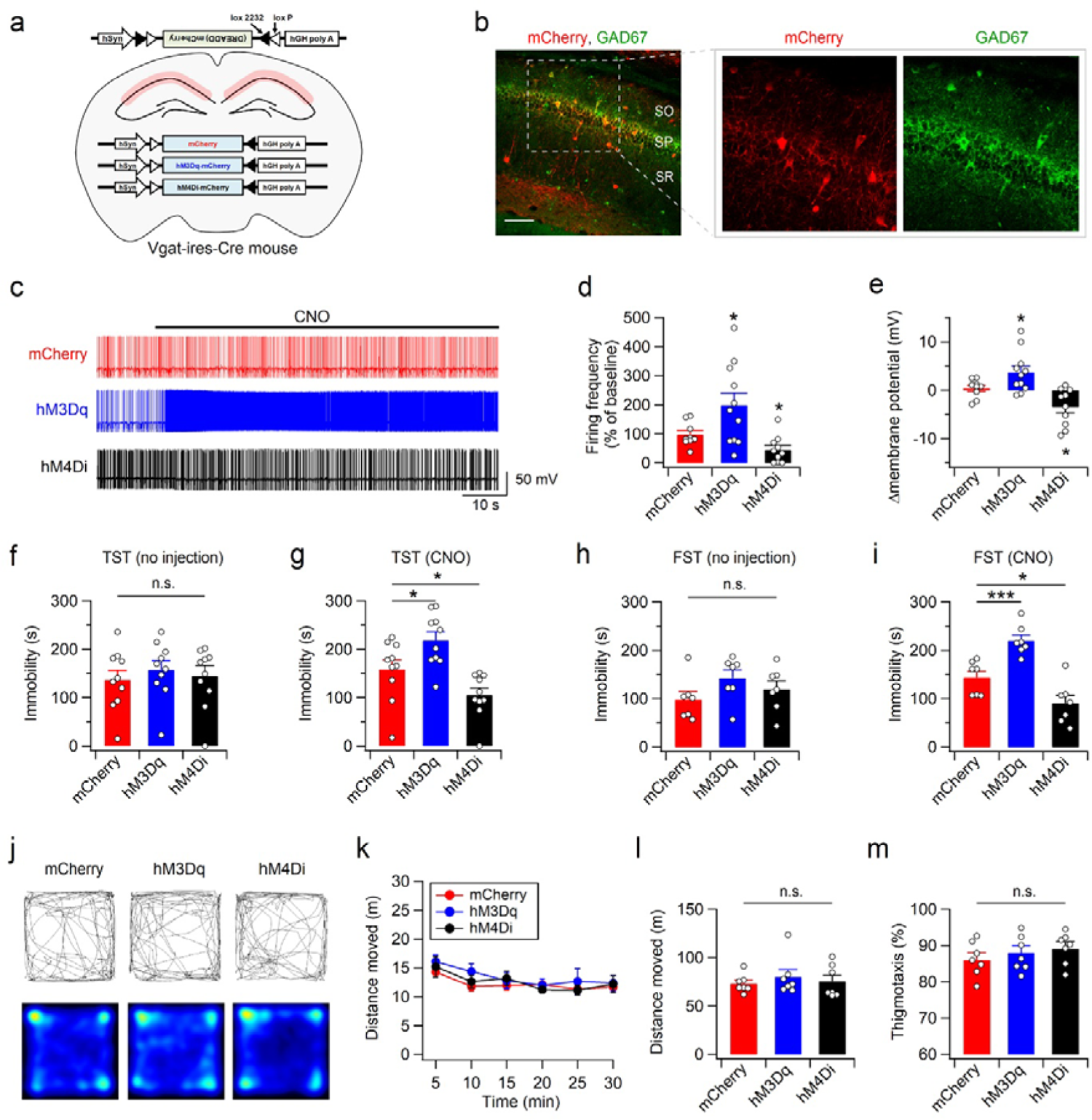
Pharmacogenetic manipulation of CA1 interneurons alters behavioral despair. (a) Design of Cre-dependent expression of mCherry, hM3Dq-mCherry or hM4Di-mCherry in the CA1 interneurons. (b) Double immunostaining of hippocampal sections with mCherry (red) and GAD67 (green) antibodies reveals interneuron-specific expression of mCherry in the CA1 subfield of Vgat-Cre mice. Scale bar, 100 µm. SO, stratum oriens; SP, stratum pyramidale; SR, stratum radiatum. (c) Representative voltage traces recorded from interneurons expressing mCherry, hM3Dq-mCherry, or hM4Di-mCherry before and during the bath application of CNO (5 µM). (d) Quantification of the CNO effects on firing rate. The firing rates under CNO are expressed as a percentage of baseline (absence of CNO) values. (e) Changes in membrane potential in response to CNO application are summarized. (f-i) Bar graphs represent total time spent immobile during the TST (f, g) and FST (h, i). In the absence of CNO, DREADD expression does not affect despair-like behaviors in the TST (f) and FST (h). CNO-induced activation or suppression of CA1 interneurons expressing DREADDs bidirectionally modifies immobility time in the TST (g) and FST (i). CNO was administered 30 min before the test. (j) Sample path recordings during the first 5 min (top) and entire 30-min period (bottom) in the open field test. (k) The open-field activities of the control (mCherry) and DREADD-expressing mice with 5-min intervals measured 30 min after CNO injection. (l, m) Quantification of the total distance moved (l) and thigmotaxis (m) in the open field box.

We confirmed the action of clozapine-N-oxide (CNO) in controlling the activity of DREADD-expressing neurons using whole-cell patch-clamp recordings. A bath application of CNO rapidly depolarized the membrane potential and significantly increased the firing rate in hM3Dq-expressing neurons, whereas hM4Di-expressing neurons exhibited hyperpolarization and decreased firing rates in response to CNO perfusion (Fig. 4c-e). In the absence of DREADDs (DiO-mCherry), CNO did not affect membrane potential or firing rate.

We next explored the role of CA1 interneuron activity in modulating behavioral despair (Fig. 4f-i). The expression of DREADDs in inhibitory neurons did not affect behavioural despair. In the absence of CNO, all groups spent a similar amount of time immobile during both the TST (Fig. 4f) and FST (Fig. 4h). Interestingly, CNO injection markedly increased the immobility time of hM3Dq-expressing mice in the TST (Fig. 4g) and FST (Fig. 4i). In contrast, hM4Di-expressing mice exhibited reduced immobility in both the TST and FST following CNO injection. However, CNO injection did not produce any statistically significant changes in the immobility time of mCherry-expressing mice (TST: t_(9)_ = −0.726, P = 0.487; FST: t_(6)_ = −2.418, P = 0.052; paired *t*-test). CNO-induced changes in TST and FST behaviors were unlikely to arise from altered general activity or anxiety, as hM3Dq- and hM4Di-expressing mice showed similar locomotor activities and thigmotaxis levels when compared with mCherry-expressing mice in the open-field test (Fig. 4j-m).

We further examined whether a specific type of hippocampal interneurons was associated with despair-like behaviors (Fig. 5). On injecting AAV2-hSyn-DIO-DREADDs-mCherry into the hippocampal CA1 area of PV-Cre knock-in (PV-Cre) mice, mCherry signals were mainly detected in PV-positive cells (Fig. 5a,b). A small portion of mCherry-expressing cells was immunostained with anti-SST antibodies, indicating DREADD expression in cells expressing both PV and SST (Supplementary Fig. 8). Infection of WT mice with a mixture of AAV-SST-Cre and AAV2-hSyn-DIO-DREADDs-mCherry predominantly expressed DREADDs-mCherry in SST-expressing cells (Fig. 5c, d and Supplementary Fig. 9). However, pharmacogenetic manipulation of both types of interneurons produced similar effects on immobility behaviors in the TST and FST (Fig. 5e-l). Activation of PV- or SST-expressing cells increased immobility behaviors, while suppression of either type of interneuron decreased despair-like behaviors. Consistent with the pharmocogenetic manipulation of Vgat-positive interenurons, altered activity in PV or SST-positive cells did not significantly impact open-field behaviors of mice (Fig. 5m-t). Collectively, these results indicate that changes in CA1 interneuron activity are sufficient for modulating despair-like behaviors.

**Fig. 5:**
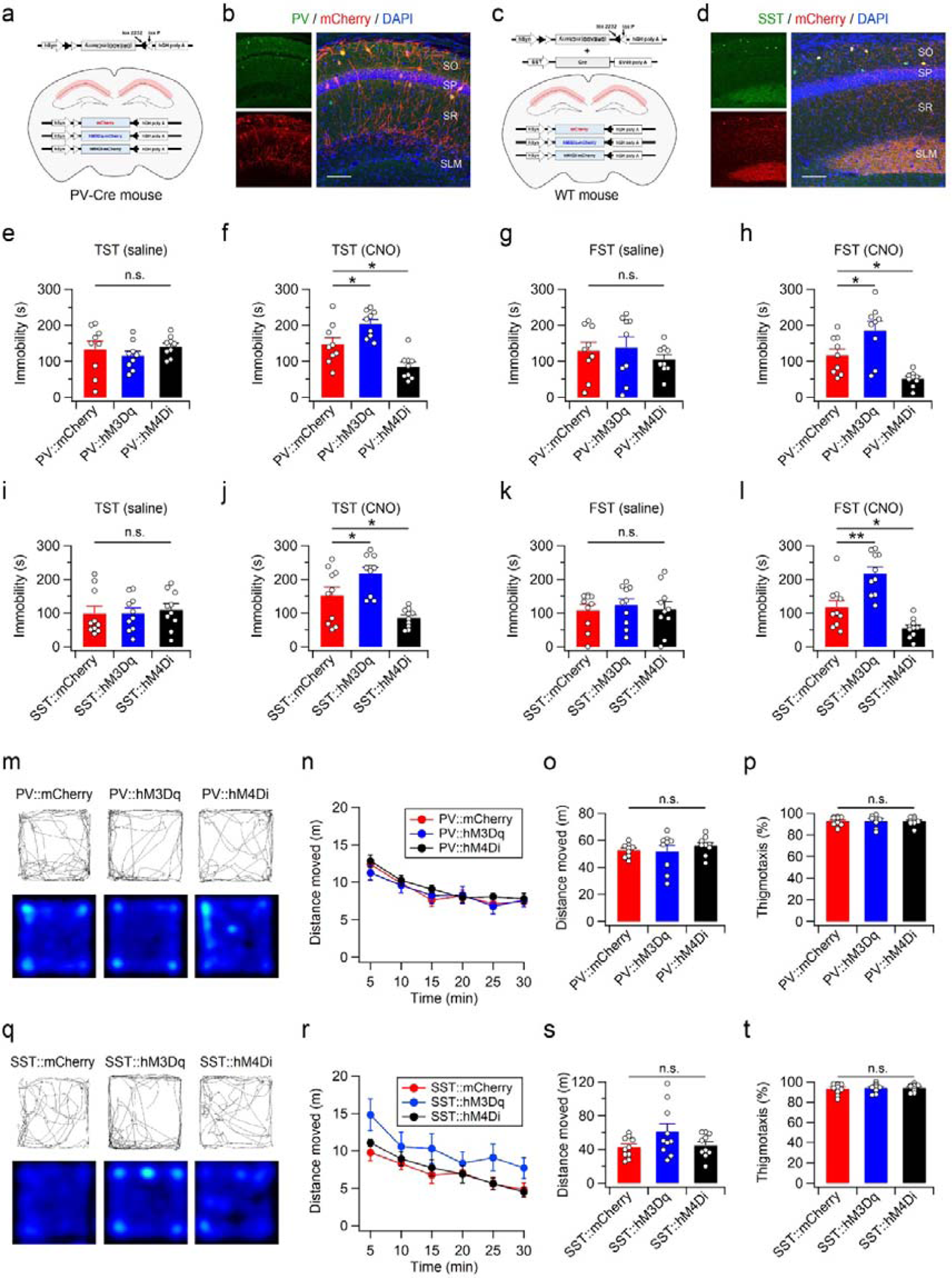
Pharmacogenetic manipulation of PV or SST interneurons in the hippocampal CA1 area is sufficient to modify behavioral despair. (a) Design of Cre-dependent expression of mCherry, hM3Dq-mCherry or hM4Di-mCherry in the PV-positive interneurons. (b) Immunohistochemical staining of hippocampal sections from PV-cre mice showing expression of mCherry in the PV-expressing cells. (c) Schematic depiction of Cre-dependent expression of mCherry, hM3Dq-mCherry or hM4Di-mCherry in the SST interneurons. (d) Expression pattern of mCherry in the hippocampal CA1 area of WT mice co-infected with AAV-DiO-DREADDs-mCherry and AAV-SST-Cre. Scale bars, 100 µm. (b, d). (e-l) Manipulation of PV or SST interneuron activity modifies immobility behaviors in the TST and FST. TST immobility time of PV-Cre mice infected with DiO-mCherry, DiO-hM3Dq-mCherry or DiO-hM4Di-mCherry was measured 30 min after saline (e) or CNO (f) injection. (g, h) FST immobility time of PV-Cre mice measured 30 min after saline (g) or CNO (h) injection. TST immobility time of mice expressiong mCherry, hM3Dq-mCherry or hM4Di-mCherry in the CA1 SST interneurons in the absence (i) or presence (j) of CNO. FST immobility time measured 30 min after saline (k) or CNO (l) injection in mice expressing mCherry or DREADD-mCherry in SST neurons. Inter-test intervals were 7-8 days (e-l). (m-p) Open-field activity of mice is not significantly affected by pharmacogenetic manipulation of PV interneurons in the CA1 area. Example path recordings of PV-Cre mice during the first 5 min (m, top) and entire 30-min period (m, bottom) in the open field test. CNO was administered 30 min before the test. Quantification of the distance moved across 5-min time bins (n), entire 30-min period (o), and thigmotaxis (p) in the open field box. (q-t) same as (m-p) but for mice expressing mCherry or DREADD-mCherry in SST neurons.

### Pharmacological disinhibition induces rapid eEF2 activation in the hippocampus and transient antidepressant-like behavior

RalBP1^-/-^ mice exhibited enhanced eEF2 activation in the hippocampus (Fig. 2), but the expression levels and function of NMDARs in RalBP1^-/-^ CA1 neurons remained unaltered^12^. Based on these observations, we hypothesized that reduced synaptic inhibition and the resultant activation of principal neurons might be sufficient to activate eEF2 signaling. To assess this hypothesis, we examined the effect of disinhibition induced by the GABA_A_R antagonist pentylenetetrazol (PTZ) on phosphorylation and expression levels of eFF2 in the WT hippocampus. Consistent with a previous pharmacological study^29^, bath application of PTZ (145 µM) rapidly reduced the amplitude of evoked IPSCs in CA1 pyramidal neurons in hippocampal slices, reflecting the disinhibition of CA1 pyramidal neurons (Fig. 6a-c). This concentration is considered to be the brain PTZ concentration (∼20 µg/mL) achieved 30 min after intravenous injection of 20 mg/kg in rodents^30^. A single administration of a subconvulsive dose (20 mg/kg, i.p.) of PTZ induced rapid (< 30min) and significant dephosphorylation of eEF2 in the hippocampus. However, the PTZ-induced activation of eEF2 did not alter BDNF levels or activation of mTOR and ERK signaling in the hippocampus (Fig. 6d).

**Fig. 6:**
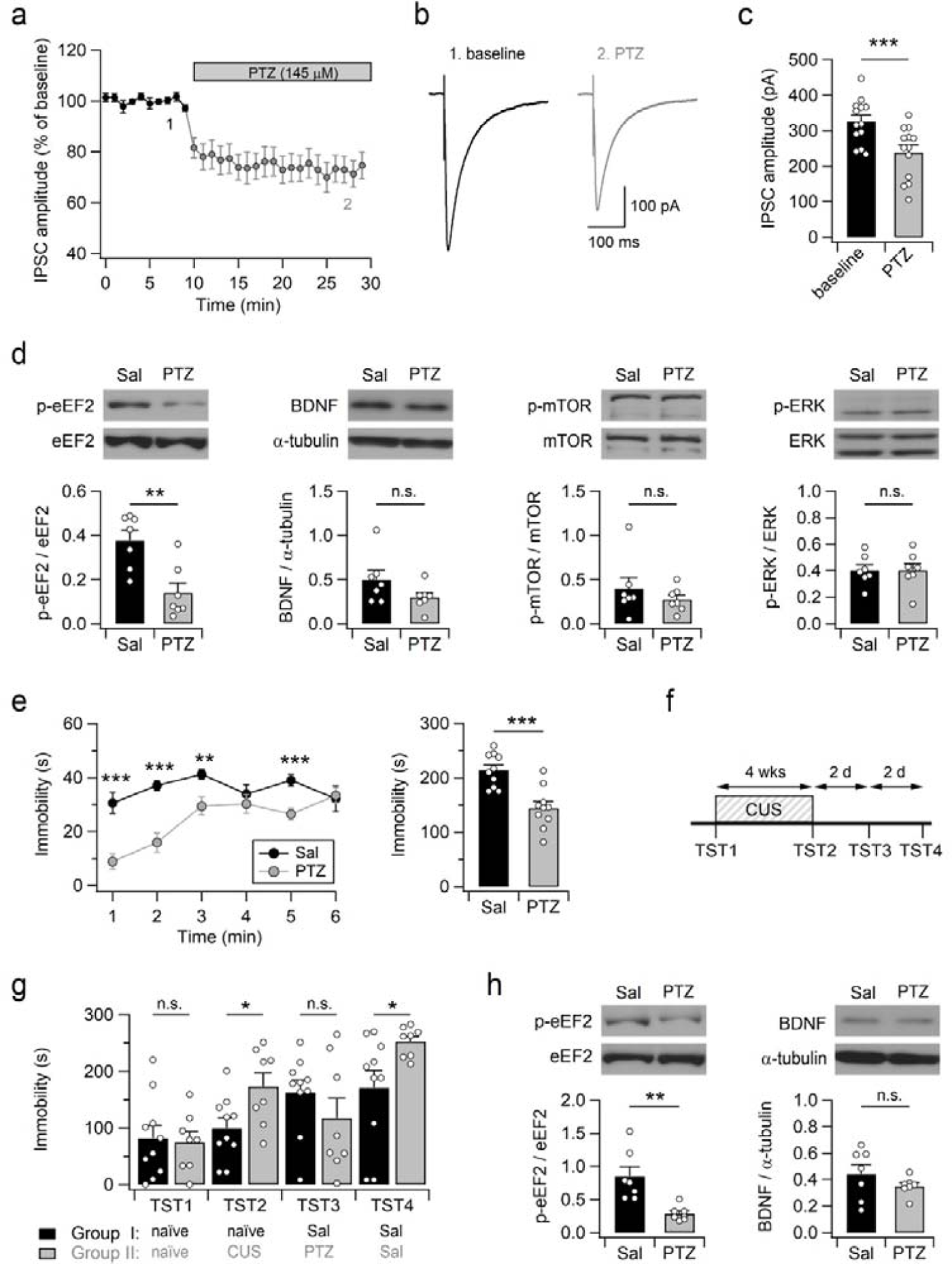
PTZ induces rapid and sustained eEF2 dephosphorylation in the hippocampus but transient antidepressant effects. (a) The peak amplitudes of evoked IPSCs recorded in the CA1 pyramidal neurons are plotted against time. (b) Sample traces of eIPSCs measured during the baseline (1, 5−10 min) and PTZ perfusion (2, 25−30 min). (c) Mean amplitudes of eIPSCs measured in the absence and presence of PTZ. (d) Representative western blots for total and phosphorylated forms of eEF2, mTOR and ERK, and BDNF in hippocampal samples obtained 30 min after treatment (top). Densitometric analysis shows the selective effect of PTZ on eEF2 but not BDNF, mTOR and ERK signaling in the hippocampus (bottom). (e) TST immobility is reduced in mice received PTZ. Bar graph represents total immobility time over 6 min (right). TST was performed 30 min after PTZ injection. (f) Experimental scheme. After scoring the immobility time of naïve WT mice in the TST (TST1), one group of mice were exposed to CUS for 4 weeks. CUS-induced despair-like behaviors were analyzed by the TST (TST2) on the next day after the end of CUS. Thirty minutes before the third TST (TST3), control and stressed mice received saline and PTZ, respectively. Both groups of mice were injected with saline 30 min before the fourth TST (TST4). (g) PTZ administration transiently reduces despair-like behaviors induced by CUS. (h) Single PTZ administration induces sustained activation of eEF2 signaling in the hippocampus. Representative western blots for eEF2 and BDNF in hippocampal lysates prepared 48 h after PTZ injection (top). Quantification of p-eEF2/eEF2 ratio and BDNF levels in the hippocampus (bottom). (d, h) Blots were cropped, and full blots are presented in Supplementary Fig. 13.

We next examined the effect of acute PTZ administration on TST behaviors. A single administration of PTZ (20 mg/kg, i.p.) significantly reduced the immobility time of mice during the 6-min TST (Fig. 6e). This concentration of PTZ is a sub-convulsant dose, as no detectable convulsions, seizure behaviors, or hyperactivity were observed (Supplementary Fig. 10). We further examined the effect of PTZ-induced disinhibition on stress-induced despair-like behaviors. Accordingly, we measured the immobility time of WT mice in the TST (TST1; Fig. 6f, g) and assigned mice to two experimental groups. One group received chronic unpredictable stress (CUS) for 4 weeks (Fig. 6f). Mice exposed to CUS spent significantly more time immobile when compared with controls, indicating CUS-induced despair-like behaviors in the stressed group (TST2; Fig. 6g). Furthermore, acute PTZ administration (20 mg/kg, i.p., 30 min before the test) effectively improved the despair-like behaviors of stressed mice, as both mice groups spent similar amounts of time immobile in the third TST (TST3; Fig. 6g). Intriguingly, a single PTZ administration did not induce a long-lasting behavioral change in stressed mice. Mice exposed to chronic stress displayed enhanced despair-like behavior when compared with control mice in the fourth TST, performed two days after PTZ administration (Fig. 6g). As PTZ has a short half-life (2.5 – 3.8 h) in rodents^31^, the rapid and short-lasting behavioral effects of PTZ could be attributed to transient synaptic disinhibition and activation of eEF2. We then wondered whether eEF2 activity would return to the basal level within 2 days of PTZ administration. Using a different cohort of unstressed mice, we investigated hippocampal eEF2 phosphorylation levels and TST behaviors of mice 48 h after drug administration. Unexpectedly, the phosphorylation levels of eEF2 were still lower in the PTZ group than those detected in the vehicle group (Fig. 6h). Moreover, despite sustained eEF2 activation, hippocampal BDNF levels did not differ between the two groups. Similar to mice exposed to CUS, antidepressant behavior induced by a single PTZ administration did not persist for 48 h in unstressed mice (Supplementary Fig. 11). These findings further indicate that the activation of BDNF signaling may be required for sustained antidepressant behaviors.

### Ketamine induces rapid activation of CA1 pyramidal neurons and anti-despair-like behaviors

In hippocampal slice electrophysiology, it has been reported that 1 µM of ketamine, a rapid-acting antidepressant, disinhibits CA1 pyramidal neurons and consequently increases the activity of these neurons^7^. However, it remains unclear whether an antidepressant dose (5 mg/kg, i.p.) of ketamine would rapidly increase hippocampal neuronal activity *in vivo*. Accordingly, we examined hippocampal c-Fos expression as an indicator of neuronal activiation. Indeed, the number of cells expressing c-Fos protein was significantly increased in both the hippocampus and mPFC within 1 h after ketamine injection (Fig. 7a,b and Supplementary Figure 12). Interestingly, in ketamine-treated mice, most c-Fos+ cells in the CA1 area were distributed in the principal cell layer and did not exhibit immunoreactivity for PV or SST (Fig. 7c,d). Considering that alterations in the hippocampal circuitry are sufficient to modify despair-like behaviors (Figs. 2-5), these results suggest that rapid enhancement in the activity of CA1 pyramidal neurons may contribute to the antidepressant effect of ketamine.

**Fig. 7:**
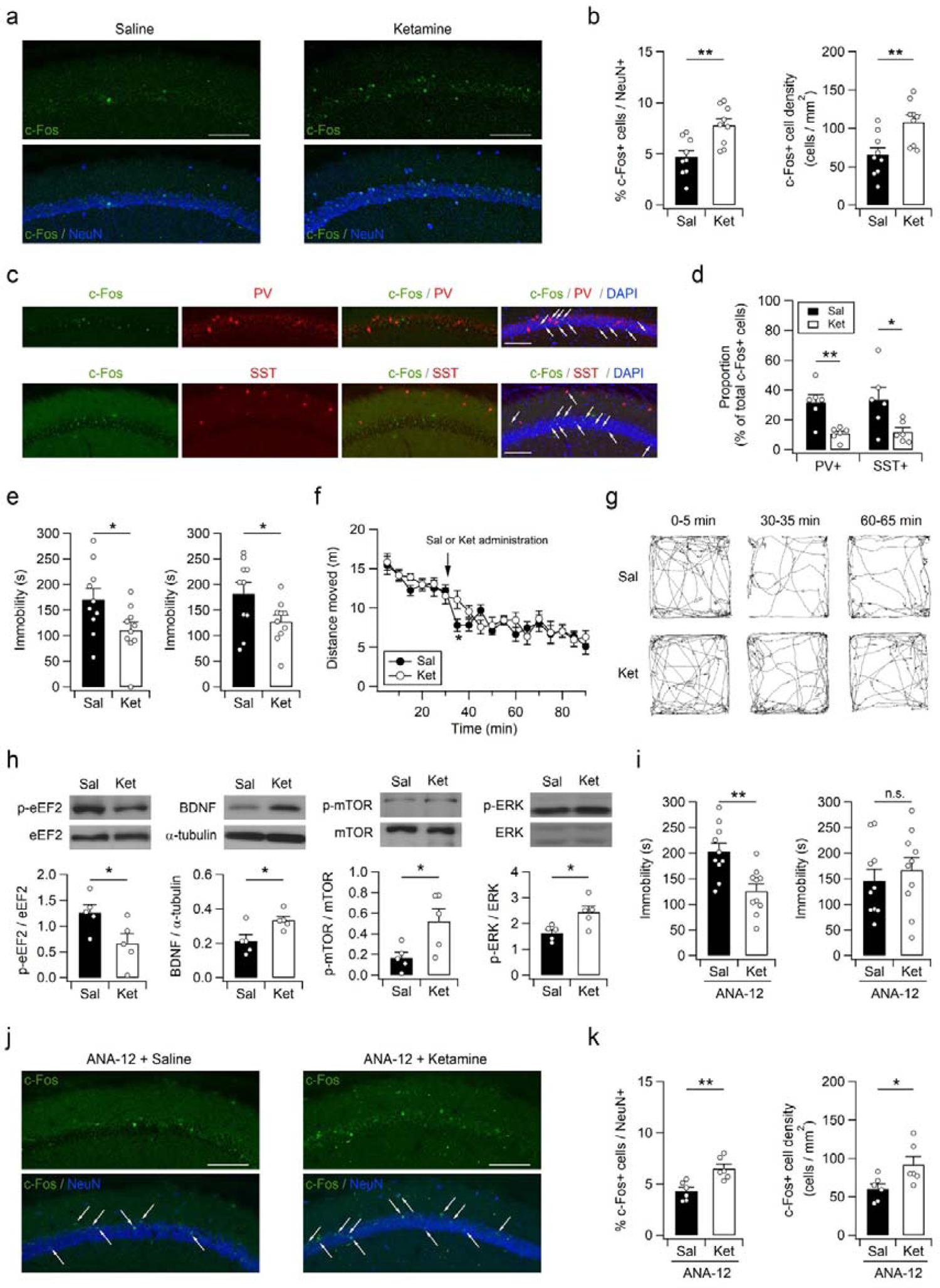
Ketamine induces rapid c-Fos expression in the hippocampus and anti-despair-like behaviors. (a) Immunohistochemical staining of hippocampal sections with the c-Fos antibody shows rapid activation of neurons by ketamine. To avoid possible effects of novel context on hippocampal neurons, mice received saline (Sal) or ketamine (Ket) and anesthetics (1 h after Sal or Ket injection) in their homecages in the vivarium. (b) Quantification of cells expressing c-Fos in the hippocampal CA1 area. (c) Hippocampal sections from ketamine-treated mice were co-stained with antibodies to c-Fos/PV (top) or c-Fos/SST (bottom). (d) Cells expressing c-Fos/PV or c-Fos/SST were measured as a percentage of total c-Fos positive cells in the hippocampal CA1 area from saline- or ketamine-treated mice. (e) Single ketamine injection induces rapid and long-lasting antidepressant effects. Reduction in TST immobility can be observed at both 30 min (left) and 48 h (right) after ketamine injection. (f) Open-field activities of mice in 5-min time bins across a 90-min session. Mice received ketamine (5 mg/kg) or saline 30 min after baseline activity recording. (g) Sample path recordings during the first 5 min, and immediately after (30-35 min) and 30 min after (60-65 min) drug administration. (h) Ketamine rapidly activates eEF2, BDNF, mTOR and ERK signaling pathways in the hippocampus. Representative western blots (top) and quantification (bottom) of proteins in hippocampal lysates obtained 30 min after ketamine injection. (i) Pretreatment (30 min before ketamine injection) with ANA-12 blocks sustained but not acute antidepressant effects of ketamine. Bar graphs represent time spent immobile during the 6 min-TST performed 30 min (left) and 48 h (right) after ketamine injection. (j) Ketamine increases the expression of c-Fos in the hippocampus in mice intraperitoneally administered ANA-12 (30 min before Sal or Ket injection). (k) Quantification of c-Fos+ cells in the hippocampal CA1 area from ANA-12-pretreated mice received saline or ketamine. Scale bars, 100 µm (a,c, and j).

Consistent with enhanced c-Fos expression, a rapid antidepressant effect was detected in the TST at 30 min after ketamine (5 mg/kg) administration (Fig. 7e). The decreased immobility during the TST at this time point was unlikely to arise from hyperlocomotion or the dissociative effects of ketamine^32^, as enhanced locomotor activities did not last for more than 10 min in the open-field test, which was performed using independent cohorts of animals (Fig. 7f,g). Notably, in contrast to PTZ, the behavioral effect of ketamine persisted for more than 48 h after a single intraperitoneal administration (Fig. 7e). We investigated the effects of ketamine on intercellular signaling 30 min after a single administration using a different cohort of mice. Ketamine exhibited similar effects to PTZ on hippocampal eEF2. However, ketamine rapidly increased the expression of the mature isoform of BDNF (m-BDNF) in the hippocampus. Moreover, phosphorylation of mTOR and ERK in the hippocampus was significantly enhanced within 30 min of ketamine administration (Fig. 7h), indicating the activation of downstream signaling pathways of BDNF. If activation of BDNF signaling is associated with the long-lasting antidepressant effects of ketamine, suppression of BDNF downstream pathways would preferentially impair the sustained effect of ketamine, leaving the transient effect intact. m-BDNF exhibits a higher affinity for the TrkB receptor than the p75 neurotrophin receptor, and the cellular effects of m-BDNF are mainly mediated via the TrkB receptor^33, 34^. The TrkB receptor antagonist ANA-12 penetrates the blood-brain barrier and reaches its peak concentration in the mouse brain as early as 30 min after intraperitoneal injection^35^. We, therefore, pretreated mice with ANA-12 (0.5 mg/kg, i.p.) 30 min before ketamine injection and assessed TST behaviors both 30 min and 48 h after ketamine administration (Fig. 7i). Ketamine still exhibited antidepressant effects in ANA-12 pretreated mice at 30 min after treatment, such that the immobility time during the TST was significantly reduced by ketamine. This finding is consistent with previous results that pretreatment (30 min) with intracerebroventricular administration of K252a, another TrkB receptor antagonist, did not block the antidepressant effect in the TST at 30 min after ketamine treatment^36^. Surprisingly, the antidepressant effects of ketamine did not persist for 48 h in ANA-12 pretreated mice (Fig. 7i). Forty-eight hours after the saline or ketamine injection, both groups displayed a similar amount of immobile time in the TST. Consistent with these observations, ketamine rapidly increased hippocampal c-Fos expression in ANA-12 pretreated (30 min before ketamine injection) mice, indicating a transient increase in neuronal activity by disinhibition (Fig. 7j,k). Collectively, these results implies that enhanced activity of hippocampal principal neurons may contribute to the rapid antidepressant effect of ketamine and that elevation of BDNF levels and the activation of downstream signalling pathways are responsible for the long-lasting antidepressant effects of ketamine^9, 37, 38^.

## Discussion

The present study suggests that alterations in synaptic inhibition in CA1 neurons affect despair-like behaviors in mice. RalBP1 mutant mice exhibited anti-despair-like behaviors and reduced synaptic inhibition with normal excitatory synaptic transmission in the hippocampus. The reduced despair-like behaviors of RalBP1^-/-^ mice were reversed by pharmacological potentiation of inhibitory synaptic transmission in CA1 neurons. Hippocampal CA1-specific suppression of inhibitory synaptic transmission via virus-mediated knockdown of Gabrg2 in WT mice recapitulated the anti-despair-like behaviors observed in RalBP1 mutant mice. Chemogenetic manipulation of CA1 interneuron activity bidirectionally modified despair-like behaviors in mice without affecting general locomotor activity and anxiety. Collectively, these results indicate that GABAergic inhibitory control of CA1 pyramidal neurons regulates behavioral despair. Furthermore, the present study provides experimental evidence suggesting that increase in the activity of CA1 pyramidal neurons may contribute to the antidepressant effects of ketamine.

It has been suggested that defects in GABAergic circuitry and an increased synaptic excitation-inhibition (E/I) ratio in the brain are associated with depressive disorders^39, 40^. Decreased GABA and increased glutamate levels have been observed in the occipital cortex in major depressive disorder (MDD)^41^, and a postmortem study showed fewer GABAergic neurons in the dorsolateral PFC in MDD^42^. Consistent with these clinical observations, heterozygous deletion of the GABA_A_R γ2 subunit (γ2^+/−^) induces depression-like behaviors in mice^28^. Interestingly, γ2^+/−^ mice exhibit a homeostatic-like reduction in the expression levels of glutamate receptors, as well as functional impairment of glutamatergic synapases in the hippocampus and mPFC. These glutamatergic defects and depression-like behaviors in γ2^+/−^ mice were reversed by ketamine^28^. In addition to GABAergic defects, glutamatergic defects such as reduced spine density in pyramidal neurons and impaired excitatory synaptic transmission and plasiticty in the hippocampus were consistently observed in stressed rodents^4, 43–47^. These alterations in excitatory and inhibitory synapses may result in abnormal neuronal activity in the hippocampus. However, the E/I ratio in the hippocampus of patient with depression remains unknown.

Based on the anatomical connectivity and deficit lesion studies, it is widely accepted that the hippocampus is functionally segregated along the hippocampal longitudinal axis; dorsal (posterior in primates) and ventral (anterior in primates) regions of the hippocampus contribute to cognitive and emotional processing, respectively^48–50^(but see^51^). Although the brain area responsible for despair remains elusive, recent studies support our observation that dorsal hippocampal network activity affects depression-related behaviors, at least despair-like behaviors in rodents. Suppression of dendritic expression of hyperpolarization-activated cyclic nucleotide-gated (HCN) channels in CA1 neurons of the dorsal hippocampus enhances neuronal excitability, while behavioral despair was reduced^52^. Restoration of HCN channel function by viral expression of TRIP8b in CA1 pyramidal neurons of the dorsal hippocampus reverses neuronal excitability and anti-despair-like behaviors in TRIP8b mutant mice^53^. Because neural network activity is determined by synaptic function and cellular excitability, it is likely that reduced synaptic inhibition and enhanced neuronal excitability similarly influence neural network activity. We observed that bilateral infusion of muscimol into the dorsal hippocampal CA1 area increased the immobility of WT mice in a dose-dependent manner and reversed the anti-despair-like behaviors of RalBP1^-/-^ mice during the TST (Fig. 2). Moreover, previous pharmacological studies have provided evidence that manipulations of the dorsal hippocampus modify despair-like behaviors in rodents. Stereotaxic infusion of UFP-101 (a nociceptin/orphanin-FQ opioid peptide receptor antagonist), 1-aminocyclopropanecarboxylic acid (a ligand for the glycine modulatory site of the NMDAR complex), CGP 37849 (a competitive NMDA receptor antagonist), imipramine (a tricyclic antidepressant), and BDNF into the dorsal hippocampus induced anti-despair-like behaviors in rodents^37, 54–56^. Thus, in addition to cognitive processing, the dorsal hippocampus seems to affect despair-like behaviors rather than anxiety or anhedonia. Considering that mPFC neurons have direct synaptic connections with axon terminals of CA1 neurons from both dorsal and ventral hippocampi^57, 58^, dorsal hippocampal projections to the mPFC might influence despair-like behaviors. Intriguingly, it has been suggested that despair consists of cognitive, emotional, behavioral, and biological domains in humans. Cognitive despair includes hopelessness, guilt, worthlessness and learned helplessness, while emotional despair comprises sadness, loneliness, anhedonia and apathy^59^. However, the despair domain of immobility behavior in rodents during the TST or FST remains uncertain and requires further investigation and characterization.

Immobile behavior in the TST and FST is believed to represent behavioral despair, by reflecting either a failure to persist in escape-directed behavior or the development of passive-coping behavior^60, 61^. Additionally, behavioral changes in these tests may result from behavioral adaptation or adaptive cognitive processes, at least in the FST^62^. Indeed, the mean immobility time tended to increase when mice were exposed to TSTs more than three times (Fig. 2c and 6g), indicating habituation or adaptive learning. However, no significant difference in the immobility time was observed in control group animals across multiple TSTs (Supplementary Table 1). Moreover, a genotype-specific phenotype was still observed in the third TST (Fig. 2c), although intrahippocampal infusion of muscimol reversed the abnormal TST behaviors of RalBP1^−/−^ mice in the previous test. Similarly, we observed bidirectional effects of CUS (TST2) and PTZ treatments (TST3) on immobile behavior, and recovery (TST4) after PTZ treatment (Fig. 6g). Furthermore, manipulation of hippocampal inhibitory circuitry produced similar effects on immobile behaviors in both the TST and FST, without affecting general locomotor activity. However, further investigation is needed to determine whether immobile behaviors in these tests are affected by coping strategies of mice to stressful stimuli or other hippocampus-dependent cognitive functions.

Our results suggest that the anti-despair-like behavior can be attributed to the increased activity of hippocampal CA1 neurons caused by disinhibition or BDNF-induced potentiation of excitatory synaptic transmission. A recent study has revealed that ketamine induces long-lasting (24 h >) enhancement of excitatory synaptic transmission in CA1 pyramidal neurons^63^. In the present study, DREADD-mediated manipulation of CA1 interneuron activity rapidly modified behavioral despair. Pharmacological suppression of inhibitory transmission by PTZ (20 mg/kg) administration was sufficient to induce rapid anti-despair-like behaviors in both control and stressed mice (Fig. 6). A previous study has shown that a low dose (1 mg/kg, i.p.) of the GABA_A_R antagonist picrotoxin did not affect the immobility behaviors of mice during the FST^9^. Consistent with this result, oral administration of 10 mg/kg, but not 2.5 and 5 mg/kg, picrotoxin produced a statistically significant reduction in the duration of immobility during the FST^64^. Furthermore, subconvulsant doses of bicuculine (a GABA_A_R antagonist) and strychnine (a glycine receptor antagonist) significantly reduce behavioral despair in mice^64^.

Importantly, however, PTZ did not exhibit sustained antidepressant effects or accompany BDNF elevation in the hippocampus. Considering the short half-life (2.5-3.8 h) of PTZ in the rodent brain^31^, it is likely that the increased E/I ratio induced by disinhibition was restored to the basal level after complete clearance of PTZ in the absence of BDNF-induced excitatory synaptic potentiation. RalBP1 mutant mice also displayed decreased despair-like behaviors and reduced levels of phosphorylated eEF2, with normal BDNF levels. Reduced interneuron density might result in permanently enhanced activity in CA1 pyramidal neurons and lead to stable anti-despair-like behavior in RalBP1^-/-^ mice. Decreased immobility of RalBP1^-/-^ and PTZ-treated mice in the TST and FST are unlikely to originate from epileptic seizures. RalBP1^−/−^ mice did not exhibit convulsive seizures, epileptic EEG rhythms, or behavioral signs of absence seizures, such as sudden behavioral arrest, nodding, whisker trembling, tail flicking, or hyperactivity. The PTZ dose used in behavioral and biochemical analyses was not sufficiently high to induce overt seizure activity in mice (Supplementary Figure 10).

Ketamine exhibited rapid (< 30 min) and sustained (∼ 48 h) behavioral effects (Fig. 7). Ketamine more effectively antagonizes NMDARs on GABAergic interneurons than on pyramidal neurons, resulting in the decreased firing of interneurons and subsequent pyramidal neuron bursts^4, 7, 65, 66^. Consistent with a previous electrophysioloy study^7^, ketamine rapidly increased c-Fos expression in the CA1 pyramidal population regardless of TrkB receptor inhibition (Fig. 7). These findings suggest that the rapid antidepressant effect of ketamine is attributable, either in full or in part, to the enhanced neuronal activity of pyramidal neurons caused by disinhibition. In contrast to mice administered PTZ, which showed transient antidepressant activity, increased BDNF levels, as well as activation of the downstream signaling molecules ERK and mTOR, were observed in the hippocampus of ketamine-treated mice. This indicates that the long-lasting antidepressant action of ketamine might originate from an enhanced E/I ratio^63^ or increased signal to noise caused by BDNF-induced neurotropic and neuroplastic changes^67–69^. Selective suppression of the sustained antidepressant effect of ketamine by the TrkB receptor antagonist ANA-12 further supports this finding (Fig. 7). Although the mechanisms underlying ketamine-induced BDNF elevation and behavioral responses are not comprehensively understood^9, 32, 70^, our results suggest that alterations in hippocampal activity contribute to the antidepressant effect of ketamine.

## Methods

### Animals

All experiments were performed using male mice (C57BL/6N), except for the EEG recordings. Animals were group-housed (3−5/cage) in a specific pathogen-free facility and maintained in a climate-controlled room with free access to food and water under a 12 h light/dark cycle. Generation and genotyping of RalBP1 mutant mice have been described previously^11, 12^. RalBP1 mutant mice and Vgat-ires-Cre (Slc32a1^tm^^2(cre)Lowl^ knock-in; Jackson Laboratory stock #016962) mice were backcrossed to C57BL/6N (Orient Bio, Sungnam, Korea) mice for at least 10 generations before use. PV-Cre knock-in (Pvalb^tm1(cre)Arbr^; Jackson Laboratory stock #017320) mice were transferred from Institute for Basic Science (IBS, Daejeon, Korea) and were backcrossed to C57BL/6N mice for 2 generations. Animal maintenance and all animal experiments were approved by the Institutional Animal Care and Use Committee (IACUC) at SNU.

### Behaviors analyses

All behavioral tests were performed between 10 a.m. and 6 p.m. with male mice that were at least 9 weeks old at the start of testing. All experimental mice were acclimated to the behavior testing room for at least 1 h prior to beginning of testing. Between trials, mazes and apparatuses were cleaned with 70% ethanol. Muscimol, pentylenetetrazol and ANA-12 were purchased from Sigma-Aldrich (St. Louis, MO, USA). Ketamine hydrochloride was purchased from Yuhan (Seoul, Korea). Most of the behavioral tests were conducted as previously described^71^, unless otherwise noted.

For tail suspension test (TST), mice were suspended by the tail from a metal hook extending from a steel bar using an adhesive tape in a 3-sided chamber with opaque walls. The distance between the floor of the chamber and the steel bar was approximately 40 cm. Mice that climbed onto their tail or fell off during the test were excluded from analysis. The duration of immobility over a 6-min session was manually scored by an experienced experimenter blind to treatment and mouse genotype.

Forced swim test (FST) was conducted by placing mice in glass beakers (20 cm high, 15 cm in diameter) containing 25−26 °C water at a 15-cm depth. An opaque plastic divider was placed between the glass beakers. Water was regularly changed and the beakers were cleaned between subjects. A mouse was judged to be immobile when it remained floating in the water without struggling or swimming and was only making minimum movements necessary to stay afloat. The total duration of immobility over the 6 min observation period was scored by an experimenter blinded to the experimental details.

Sucrose preference test was performed once daily for 3 days. Mice were deprived of water but not food for 12 h before each test and allowed to freely access to both water and 1% sucrose in individual cages for 12 h. The position of each bottle was switched daily to avoid false sucrose preference resulting from side bias. Volume of liquid consumed was determined by weighting the bottles. The preference (%) for sucrose solution was calculated as follows: (sucrose solution intake /total liquid intake) × 100%.

Open field test was conducted by placing in the center of an open field apparatus with opaque walls (40 × 40 × 40 cm) in a dimly lit room. The behavior of each mouse was monitored by video recording. The total distance traveled and time spent in the entire open field and in the center (20 × 20 cm) were calculated using video tracking software (Ethovision XT, Noldus, Netherlands).

Elevated plus maze test was performed using a plus-shaped maze with two open arms and closed arms surrounded by opaque walls (20 cm in height). The maze was elevated 50 cm from the floor, and dimension of each arm was 5 cm wide and 30 cm in length. Each mouse was place in the center of the maze and movement of the mouse was video recorded for 5 min.

The elevated zero maze consisted of black acrylic in a circular track 5 cm wide, 60 cm in diameter, and elevated 50 cm from the floor. The maze was divided into four equal quadrants, two of which were enclosed with black acrylic walls (20 cm in height). Each mouse was placed in the center of one of the two closed quadrants and movement of the mouse was video recorded for 5 min.

For chronic unpredictable stress (CUS), mice were exposed in a random order to a variety of chronic stressors, including a wet cage (12 h), light-dark cycle reversal (24 h), white noise (100 dB, 12 h), cold water swim (10 °C, 1 h), restraint (2 h), cage shake (30 rpm, 12 h) and electric foot shocks (10 scrambled shocks with duration of 2 s over 120 min). Animals were subjected to one stressor daily for 28 days.

The Morris water maze test was performed using a white circular pool (120 cm in diameter) filled with 23°C–25°C water, which was made opaque with nontoxic white paint. A transparent circular platform (10 cm in diameter) submerged 1 cm beneath the surface. Two-day pre-training sessions consisted of one trial per day, and each mouse was allowed to remain on the visible platform for 30 s. The mice were then trained to find the hidden platform for 7 days with four trials per day and inter-trial intervals of 1 min. In each trial, mice were released at one of the three starting points and allowed to find a submerged platform. If mice did not locate the platform within 90 s, they were manually placed on the platform by the experimenter. Mice remained on the platform for 30 s before subsequent trials or before being returned to home cages. The probe test was performed 1 d after the completion of the training sessions. Swim paths and quadrant occupancies of each mouse were analyzed using video tracking software (Ethovision XT, Noldus).

The accelerating rotarod test was performed using the Rotamex 5 equipment (Columbus Instruments, OH, USA). Mice were placed on motionless rods, which were then accelerated from 5 to 40 rpm over 5 min. The latencies to falling off the rotating rod were measured. Each mouse was trained once daily for 3 days.

Training sessions for contextual and cued fear conditioning consisted of a 300 s exploration period (Pre-CS, contextual), followed by the presentation of an 18 s tone and a 2 s tone paired with a foot shock (0.7 mA) in the fear conditioning chamber (Coulbourn Instruments). Afterwards, each mouse was kept in the chamber for 60 s before being returned to its home cage. Contextual fear conditioning tests were performed in the same chamber (CS, contextual) 24 h after the training by monitoring mouse activity for 5 min. For cued fear conditioning, mice were placed in a distinct chamber, and their activities (pre-CS, cued) were monitored for 180 s, followed by presentation of a 180 s tone (CS, cued). The time spent freezing was scored post hoc by an experimenter blind to genotype.

Marble burying test was performed by placing animals individually in a transparent acrylic cage containing 15 clean glass marbles (1.5 cm in diameter) spaced evenly (3 × 5) on the surface of the 5-cm-deep bedding. Animals were allowed to explore the marbles for 10 min, and the number of marbles buried with bedding up to two-thirds of their surface area was counted.

Social behavior of animals was tested using the three-chambered social approach apparatus. The apparatus was white open-topped box (60 × 40 × 20 cm) and divided into three chambers (a 20-cm-wide center chamber and 20-cm-wide side chambers) with two acrylic walls. Each of the two side chambers contained an empty wire cup. Dividing walls had retractable doorways that allowed access into each chamber. During the habituation phase, animals were placed in the center chamber and allowed to explore freely for 10 min. During the sociability phase, the animal was gently guided to the center chamber, and the two doors were blocked while a stranger C57BL/6 mouse (stranger 1) placed inside one of the wire cups and an inanimate object was placed inside the other wire cup. Then, the two doors were opened and the subject animal was allowed to explore all the three chambers for 10 min. The amount of time spent in each chamber and time spent exploring mice or objects were measured. For the social novelty test, the subject mice again gently guided to the center chamber, and the two doors were blocked while an inanimate object was replaced by a novel unfamiliar mouse (stranger 2). Afterwards, the two doors were opened and the subject animal again allowed to explore all three chambers for 10 min. The behavior of the animals was recorded and analyzed using video tracking software (Ethovision XT, Noldus). All stranger mice were males of the same age and previously habituated to being enclosed in wire cups in the apparatus for 30 min one day before the test.

Pre-pulse inhibition (PPI) test was performed using the SR-Lab Startle Response system (San Diego Instruments, Ca, USA). Mice were placed into a small transparent tube mounted on a movement sensor platform in a sound-attenuating chamber and acclimated for 10 min before the testing session. The startle pulse was 120 dB (40 ms) and pre-pulses were either 75 or 80 dB (20 ms). During the acclimation period and each test session, background white noise (65 dB) was continuously presented. Baseline startle responses of animals were measured by 5−10 startle pulse (120 dB) alone trials, with inter-trial interval of 30 s. The peak startle amplitude was determined by measuring the largest positive and negative peaks of the startle response within a 300 ms window after the onset of the acoustic stimulus. During the PPI sessions, the same startle pulse preceded (100 ms) by a 75 dB or 80 dB pre-pulse in a pseudorandom order, with 5 testing trails per each pre-pulse. Scores for each measure were averaged, and PPI was calculated as a percent reduction of startle response: PPI (%) = (startle pulse alone – startle pulse with pre-pulse) / startle pulse alone) × 100.

### Constructs and virus preparation

For Gabrg2 knockdown assay, Gabrg2 cDNA was amplified from mouse hippocampal cDNA and was cloned into pGW1 vector. Four independent shRNA (#1: gcattggaagctcagtctactctcctgta; #2: tggaatgatggtcgagttctctacacctt; #3: ccaaggtctcctatgtcacagcaatggat; #4: tggattgaggaatacaactgaagtagtga) constructs in the pGFP-V-RS vector and a non-targeting (NT: gcactaccagagctaactcagatagtact) pGFP-V-RS plasmid were purchased from OriGene (Cat. # TL515175). For lentivirus-mediated expression, the cassettes containing the human U6 (hU6) promoter-shRNA#3 and hU6-NT were amplified by PCR, and then subcloned into the pLVX-DsRed-Monomer-C1 (Clontech) vector, displacing the CMV promoter-DsRed-Monomer cassette. The PGK promoter-puromycin resistance gene cassette in the vector was further replaced by the CMV-eGFP cassette from the pEGFP-C1 vector (Clontech). The sequence-verified plasmid was transfected into Lenti-X 293T cells (Takara Bio) together with the packaging plasmid psPAX2 (Addgene) and the envelope plasmid pMD2.G (Addgene) using a polyethylenimine (PEI)-based transfection protocol (total DNA to PEI µg ratio of 1: 2). The ratio of pLVX-shRNA: psPAXs: pMD2.G was 4:1:2 (total 300 µg / 245 mm square dish). Lenti-X 293T cells were cultivated in the high glucose (4.5 g/L) Dulbecco’s Modified Eagle’s Medium (DMEM) containing 4 mM L-glutamine, 3.7 g/L sodium bicarbonate, 10% (v/v) tetracycline-free fetal bovine serum (FBS), 1 mM sodium pyruvate, and 1% (v/v) penicillin/streptomycin. The medium was replaced with fresh medium 24 h after transfection. Lentivirus-containing medium was harvested 72 h after transfection and was briefly centrifuged at 900 g for 10 min to remove debris. Supernatant was filtered (0.45-μm pore size) for sterilization, and lentivirus particles were concentrated by 2 rounds of ultracentrifugation (10,000 g for 2 h). Pellets were resuspended in PBS, aliquoted and stored at −80 °C. Viral titer (10^12−13^ IU/mL) was determined by transfecting serially diluted virus stocks into HEK293T cells and analyzing the number of GFP-positive cells using a fluorescence-activated cell sorter (FACS).

AAV2/hSyn-DIO-mCherry (#50459), AAV2/hSyn-DIO-hM3Dq-mCherry (#44361) and AAV2/hSyn-DIO-hM4Di-mCherry (# 44362) were obtained from Addgene (Watertown, MA, USA). AA2/SST-Cre (#CV17210-AV2) was purchased from Vigene Biosciences (Rockville, MD, USA).

### Surgery and stereotaxic injection

Under deep anesthesia with a mixture of Zoletil (50 mg/kg, i.p) and Xyalazine (1 mg/kg, i.p), mice were placed in a stereotaxic device. The skin was cut over the midline and craniotomies were performed bilaterally over the hippocampus. Either purified AAV (0.5 µL/side) or lentivirus (1 µL/side) was injected into the CA1 region of the dorsal medial hippocampus (−1.9 anteroposterior, ±1.5 mediolateral, −1.8 dorsoventral from the Bregma) using a Hamilton syringe at a rate of 100 nL/min. After completion of injection, the needle (33 gauge) was remained in place for an additional 10 min to allow the diffusion of the injection medium, and then carefully retracted to prevent backflow. Experiments were performed 3−4 weeks after viral injections.

For intrahippocampal infusion of muscimol, cannula assemblies consisted of a stainless-steel guide cannula (26 gauge with 2 mm) and a stylet were bilaterally implanted into the dorsal medial hippocampus (−1.9 anterior/posterior, ±1.5 medial/lateral, −1.2 dorsal/ventral) and were affixed to the skull with dental cement. Following surgery, mice were allowed to recover for at least 10 days before microinjection. Intrahippocampal infusion (0.5 µL/side) was made using a syringe pump (Harvard Apparatus, Pump 11) at a rate of 100 nL/min. Stainless steel injection cannulas connected to the injection syringe by polyethylene tubing were inserted into the guide cannulas. Following microinjection, the injection cannulas were remained in place for 5 min to allow diffusion of the solution into the tissue and then replaced by stylets.

### Electrophysiology

Parasagittal hippocampal slices (400 µm thick) were prepared using a vibratome (Leica, Germany) in ice-cold dissection buffer (sucrose 230 mM; NaHCO_3_ 25 mM; KCl 2.5 mM; NaH_2_PO_4_ 1.25 mM; D-glucose 10 mM; Na-ascorbate 1.3 mM; MgCl_2_ 3 mM; CaCl_2_ 0.5 mM, pH 7.4 with 95% O_2_/5% CO_2_). Immediately after sectioning, the CA3 region was surgically isolated. The slices were allowed to recover at 36 °C for 1 h in normal artificial cerebrospinal fluid (ACSF: NaCl 125 mM; NaHCO_3_ 25 mM; KCl 2.5 mM; NaH_2_PO_4_ 1.25 mM; D-glucose 10 mM; MgCl_2_ 1.3 mM; CaCl_2_ 2.5 mM, pH 7.4 with 95% O_2_/5% CO_2_), and then maintained at room temperature.

All electrophysiological recordings were performed in a submerged recording chamber, which was perfused with heated (29–30°C) ACSF. The signals were filtered at 2.8 kHz and digitized at 10 kHz using a MultiClamp 700B amplifier and a Digidata 1440A interface (Molecular Devices, CA, USA). Data were analyzed by using custom macros written in Igor Pro (WaveMetrics, OR, USA).

Miniature EPSCs (mEPSCs) were measured at –70 mV with a pipette solution containing (in mM) 110 K-gluconate, 20 KCl, 8 NaCl, 10 HEPES, 0.5 QX-314-Cl, 4 Mg-ATP, 0.3 Na-GTP, and 10 BAPTA, adjusted to pH 7.25 and 290 mOsm/kg. Spontaneous action potentials and IPSCs were blocked by tetrodotoxin (TTX, 1 µM) and picrotoxin (50 µM), respectively. For miniature IPSC (mIPSC) recording, K-gluconate in the pipette solution was replaced by equimolar KCl, and currents were measured at –70 mV in the presence of TTX (1 µM), NBQX (10 µM), and AP-5 (50 µM) in ACSF.

To measure evoked EPSC/IPSC ratio (Figs. 1 and 3), whole-cell voltage clamp recordings were made using patch pipettes (3–4 MΩ) filled with solution containing (in mM) 130 CsMeSO4, 10 TEA-Cl, 10 HEPES, 4 Mg-ATP, 0.3 Na-GTP, 5 QX-314-Cl and 10 EGTA, adjusted to pH 7.25 and 290 mOsm/kg. The synaptic responses were evoked at 0.05 Hz with an ACSF-filled broken glass pipette (0.3–0.5 MΩ) placed in the proximal region of the stratum radiatum. The mean AMPAR-mediated EPSCs were obtained by averaging 30–40 traces recorded at –57 mV. Stimulation intensity was adjusted to yield a 100–300 pA EPSC peak amplitude. To isolate the GABA_A_R-mediated currents, 30–40 traces of synaptic currents were recorded at +3 mV. To examine the effect of PTZ on inhibitory synaptic transmission (Fig. 6), evoked IPSCs were measured at a holding potential of –70 mV with the same pipette solution used for the measurement of mEPSCs. NBQX (10 µM) and AP-5 (50 µM) were added to the ACSF throughout recordings to block AMPAR- and NMDAR-mediated EPSCs, respectively. The series resistance and seal resistance were monitored, and data were discarded if they changed by more than 20% during recordings.

The membrane potentials were recorded under whole-cell current clamp mode with a pipette solution containing (in mM) 110 K-gluconate, 20 KCl, 8 NaCl, 10 HEPES, 4 Mg-ATP, 0.3 Na-GTP, and 0.5 EGTA, adjusted to pH 7.25 and 290 mOsm/kg. Neurons displaying an unstable resting potential at the beginning or during the recording were discarded.

All reagents were purchased from Sigma-Aldrich (MO, USA), except for QX-314-Cl, CNO, NBQX, AP-5 from Tocris (Bristol, UK).

### Immunohistochemistry and western blotting

Immunohistochemistry and western blotting analyses were performed as described previously^71^. Briefly, mice were deeply anesthetized with diethyl ether and transcardially perfused with heparinized (10 U/mL) phosphate-buffered saline (PBS), followed by PBS-buffered 4% (w/v) paraformaldehyde (PFA). Brains were removed, post-fixed in 4% PFA for 48 h at 4 °C and cut into 60 μm coronal sections using a vibratome (VT1200S, Leica, Germany). The sections were post-fixed (1 h), permeabilized with 0.3% (v/v) Triton X-100 in PBS, incubated in blocking buffer (5% normal goat serum, 5% horse serum, 5% donkey serum, and 0.5% BSA in PBS) for 2 h. Sections were successively incubated with primary [anti-PV: Swant, (Cat. # PV27) and Millipore (Cat. # MAB1572); anti-SST: Santa Cruz Biotechnology (Cat. # sc-55565); anti-VIP: Immunostar (Cat. # 20077); anti-CCK: Abcam (Cat. # ab37274); anti-GAD67: Millipore (Cat. # MAB5406); anti-mCherry: Abcam (Cat. # ab167453); anti-GFP: Synaptic Systems (Cat. # 132 004); anti-c-Fos: Cell Signaling Technology (Cat. # 2250S); anti-NeuN: Millipore (Cat. # ABN78 and Cat. # MAB377); overnight at 4 °C] and fluorescence (Cy3, Alexa fluor 647 or FITC: Jackson ImmunoResearch Laboratories, PA, USA) conjugated-secondary (3 h at room temperature) antibodies. Between each step, the sections were rinsed 3 times for 10 min with PBS. Images were acquired using an FV 3000 confocal laser scanning microscope and processed with FV31S-SW Viewer (Olympus, Japan). Wide-field images of entire brain sections were acquired using a TCS SP8 confocal microscope and Leica Application Suite X (Leica, Germany).

For western blotting, mouse hippocampi were homogenized in buffer (320 mM sucrose, 10 mM Tris-HCl, 5 mM EDTA, pH 7.4) containing phosphatase inhibitor cocktail (GenDEPOT, TX, USA, Cat. # P3200) and proteinase inhibitor cocktail (Sigma-Aldrich, MO, USA, Cat. # P8340). Proteins were separated by SDS-PAGE and transferred to nitrocellulose membranes. The membranes were successively incubated with primary and horseradish peroxidase (HRP)-conjugated secondary (Jackson ImmunoResearch Laboratories, PA, USA) antibodies. Signals were detected by enhanced chemiluminescence (GE Healthcare, UK) and quantified using MetaMorph software (Molecular Devices, CA, USA). The following primary antibodies were obtained from commercial suppliers: anti-ERK (Cat. # 9102S), anti-p-ERK (Cat. # 9106S), anti-eEF2 (Cat. # 2332S), anti-p-eEF2 (Cat. # 2331S), anti-mTOR (Cat. # 2972S), anti-p-mTOR (Cat. # 2971S) from Cell Signaling Technology; BDNF (Cat. # ab108319) from Abcam; Gabrg2 (Cat. # 224 003) from Synaptic Systems; tGFP (Cat. # TA150041) from Origene; α-tubulin (Cat. # T5168) from Sigma-Aldrich.

### Electroencephalogram (EEG) recording

EEG surgeries and recordings were performed as previously described^71^. Mice (7-9 weeks old) were anesthesized with 1.5% isoflurane. The skin was incised over the midline, and four epidural electrodes were bilaterally anchored into the frontal and temporal regions of the skull (frontal: +2.0 mm anteroposterior, ±1.0 mm mediolateral; temporal: −2.3 mm anteroposterior, ±2.0 mm mediolateral). A reference electrode was placed in the occipital region of the skull (−4.2 mm anteroposterior). All connectors were fixed to the skull using dental cement. Mice were allowed to recover from surgery for 7–10 days. EEG signals and video recordings were simultaneously acquired for 30 min. Mice of both sexes (+/+: three males and two females; −/−: four males and two females) were used for EEG recordings, and sex differences were not observed. EEG signals were amplified (Digital Lynx acquisition system and Cheetah software, Neuralynx) and digitized at a sampling rate of 32 kHz. The data were analyzed using Neuroexplorer (Nex Technologies). Representative spectrograms and traces were obtained using Neuroexplorer and custom macros written in Igor Pro (WaveMetrics, OR, USA), respectively.

### Statistical analysis

Statistical analyses were performed using Igor Pro (WaveMetrics) and SPSS (IBM, Armonk, USA). The normality of the collected data was determined using the Shapiro-Wilk test. The Mann-Whitney test was used to compare non-normally distributed samples. Samples that satisfied normal distribution were compared using two-tailed Student’s *t*-tests. For multiple groups, one-way ANOVA, followed by with Tukey’s HSD (Honestly Significant Difference) post-hoc test was used to compare the samples. All bar graphs in the figures show the mean ± standard error of the mean (SEM). The levels of significance are indicated as follows: *P < 0.05, **P < 0.01, ***P < 0.001, n.s., not significant (p ≥ 0.05). The number of cells, slices, or animals used for each experiment and statistical analyses are provided in Supplementary Table 1.

## Data availability

All raw data supporting the findings of this study are available upon reasonable request. Source data are provided with this paper.

## Code availability

Custom Igor Pro functions for analyzing electrophysiology data within this manuscript are available upon request.

## Acknowledgements

S.H. Yoon, W.S. Song, and S.P. Oh are postgraduate students supported by Brain Korea 21 Plus Program. We thank the late Dr. Ronald S. Duman for his insightful comments and helpful suggestions on earlier drafts of the manuscript. M.-H. Kim was inspired and stimulated by the outstanding scientist during his sabbatical at Yale, 2018-2019. This study was supported by the National Research Foundation of Korea (NRF) grant funded by the Korea government (NRF-2017R1D1A1B03032935, 2017M3C7A1029609, 2018R1A5A2025964, 2020R1A2C2012103, and 2020R1A2C1014372), and by Creative-Pioneering Researchers Program through Seoul National University (SNU).

## Author contributions

S.H.Y and M.-H.K conceived the project, and designed and analyzed experiments. S.H.Y, S.W.S, and S.P.O performed experiments and analyzed data. G.C and S.J.K conducted the cannula infusion experiments. J.K performed EEG recording and analysis. S.H.Y and M.-H.K wrote the manuscript, with input from all other authors.

## Competing interests

The authors declare no competing interests.

## Notes

### Competing Interest Statement

The authors have declared no competing interest.

### Summary of Updates

Figures and Supplementary Figures updated

