## Supplementary Figures for "Altered GABAergic inhibition in CA1 pyramidal neurons modifies despair-like behavior in mice"

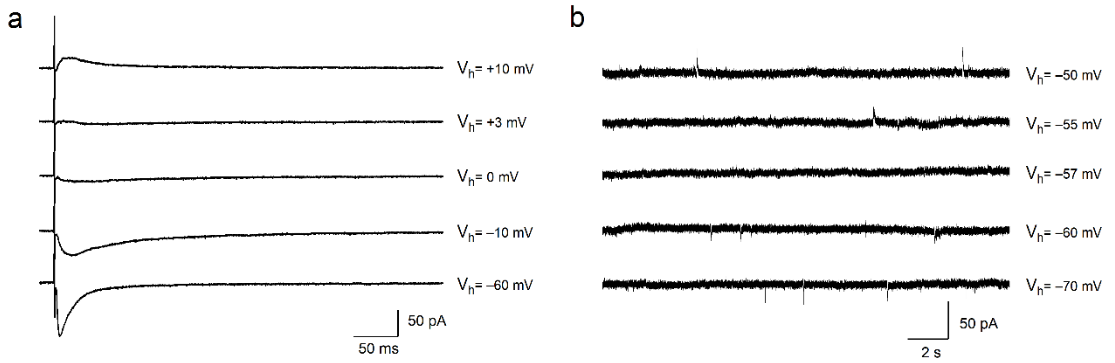


**Supplementary Fig. 1: Determination of the reversal potentials for EPSCs and IPSCs in CA1 pyramidal neurons.** (a) Sample traces of evoked EPSCs in the presence of the $GABA_AR$ blocker picrotoxin in the bathing solution. The holding potentials ($V_h$) are shown with each trace. Recording at +3 mV leads to negligible evoked EPSCs owing to the absence of a driving force for cations. (b) Example traces of spontaneous IPSCs recorded from CA1 pyramidal neurons in the presence of the AMPAR blocker NBQX and NMDAR blocker AP-5. The reversal potential of spontaneous IPSCs was −57 mV.

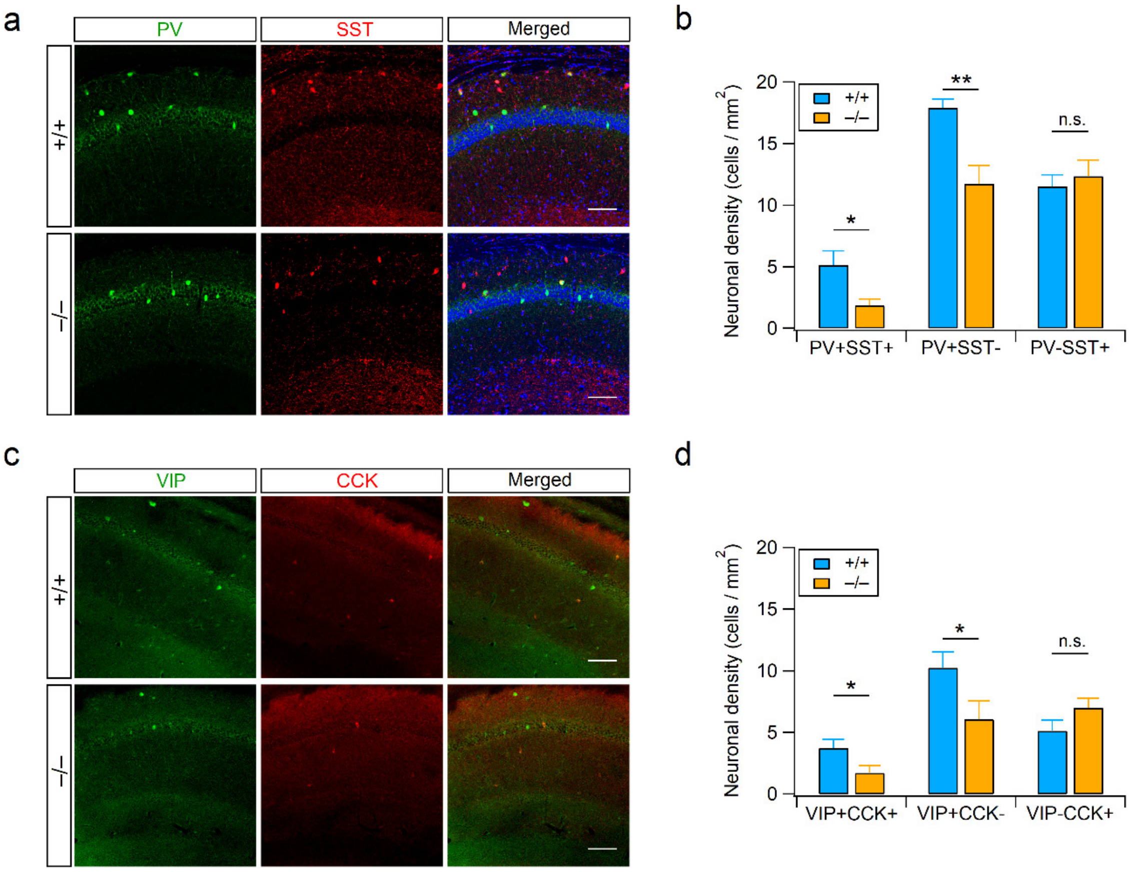


**Supplementary Fig. 2: RalBP1$^{-/-}$ mice exhibit fewer interneurons in the hippocampus.** (a) Immunohistochemical staining for parvalbumin (PV) and somatostatin (SST) in the WT and RalBP1$^{-/-}$ hippocampal CA1 areas. DAPI (blue) was used to visualize cell nuclei (right). (b) Quantification of PV- and/or SST-expressing neurons in the hippocampal CA1 area. (c) Immunostainings for vasoactive intestinal peptide (VIP) and cholecystokinin (CCK) are shown. (d) Quantification of VIP- and/or CCK-expressing neurons in the hippocampal CA1 area. (a and c) Scale bars, 100 µm.

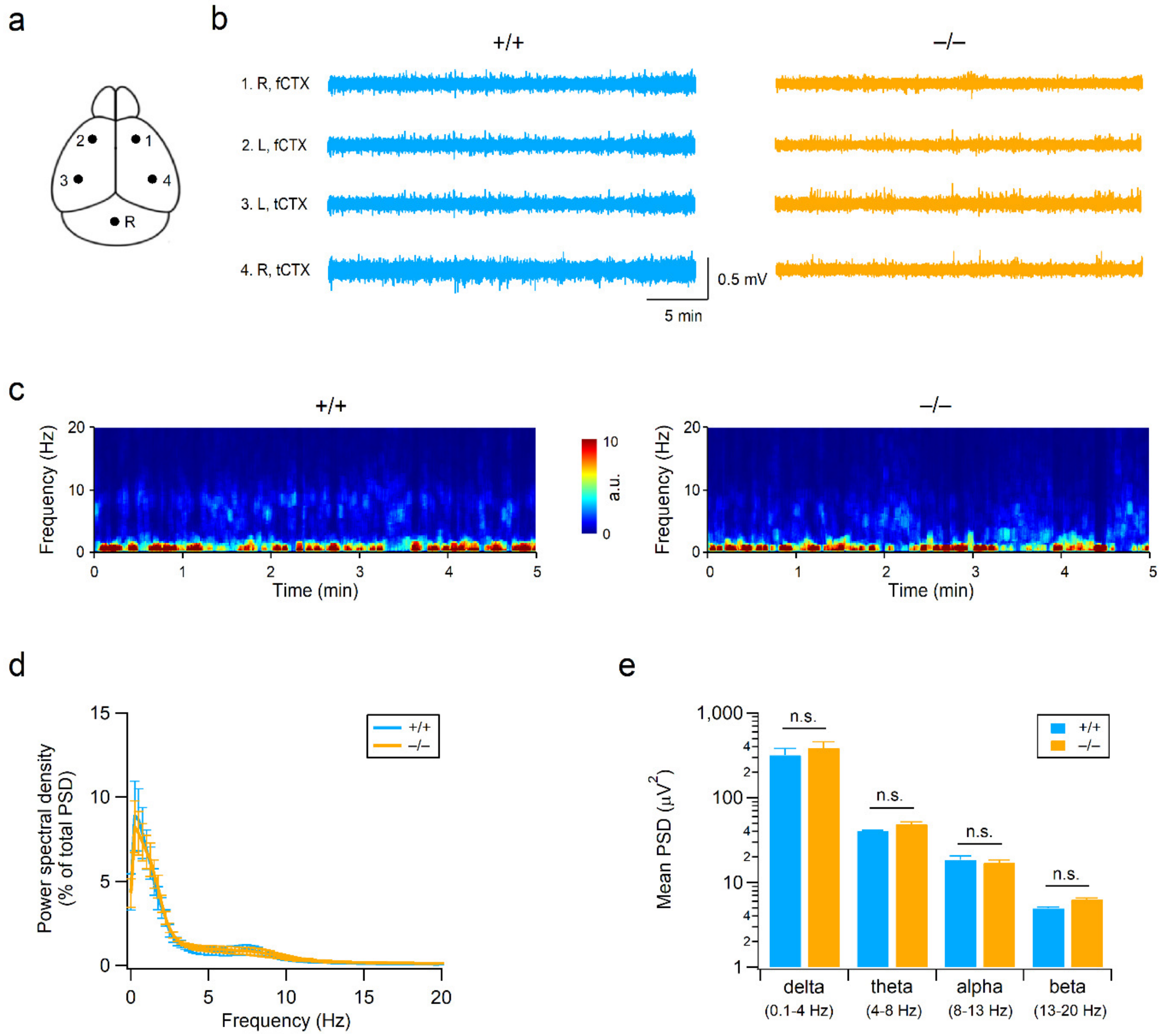


**Supplementary Fig. 3: Normal electroencephalogram (EEG) rhythms in RalBP1$^{-/-}$ mice.** (a) Diagram showing the placement of epidural electrodes in the mouse brain. R, reference electrode. (b) Sample EEG traces of RalBP1$^{+/+}$ (left) and RalBP1$^{-/-}$ (right) mice recorded for 30 min from four epidural electrodes in their homecages. No epileptic activity was observed in both genotypes. R, right; L, left; fCTX, frontal cortex; tCTX, temporal cortex. (c) Time-frequency plots of EEG waves recorded in the right temporal region of the control (left) and RalBP1$^{-/-}$ (right) brain during the first 5 min of the EEG recording. (d) Frequency characteristics of EEG traces recorded for 30 min in the right temporal region of RalBP1$^{+/+}$ and RalBP1$^{-/-}$ mice were calculated by fast Fourier transformation. (e) The average power spectral density of delta, theta, alpha, and beta rhythms calculated from RalBP1$^{+/+}$ (N = 5) and RalBP1$^{-/-}$ (N = 6) mice. n.s., not significant, $P > 0.05$ by Student's t-test.

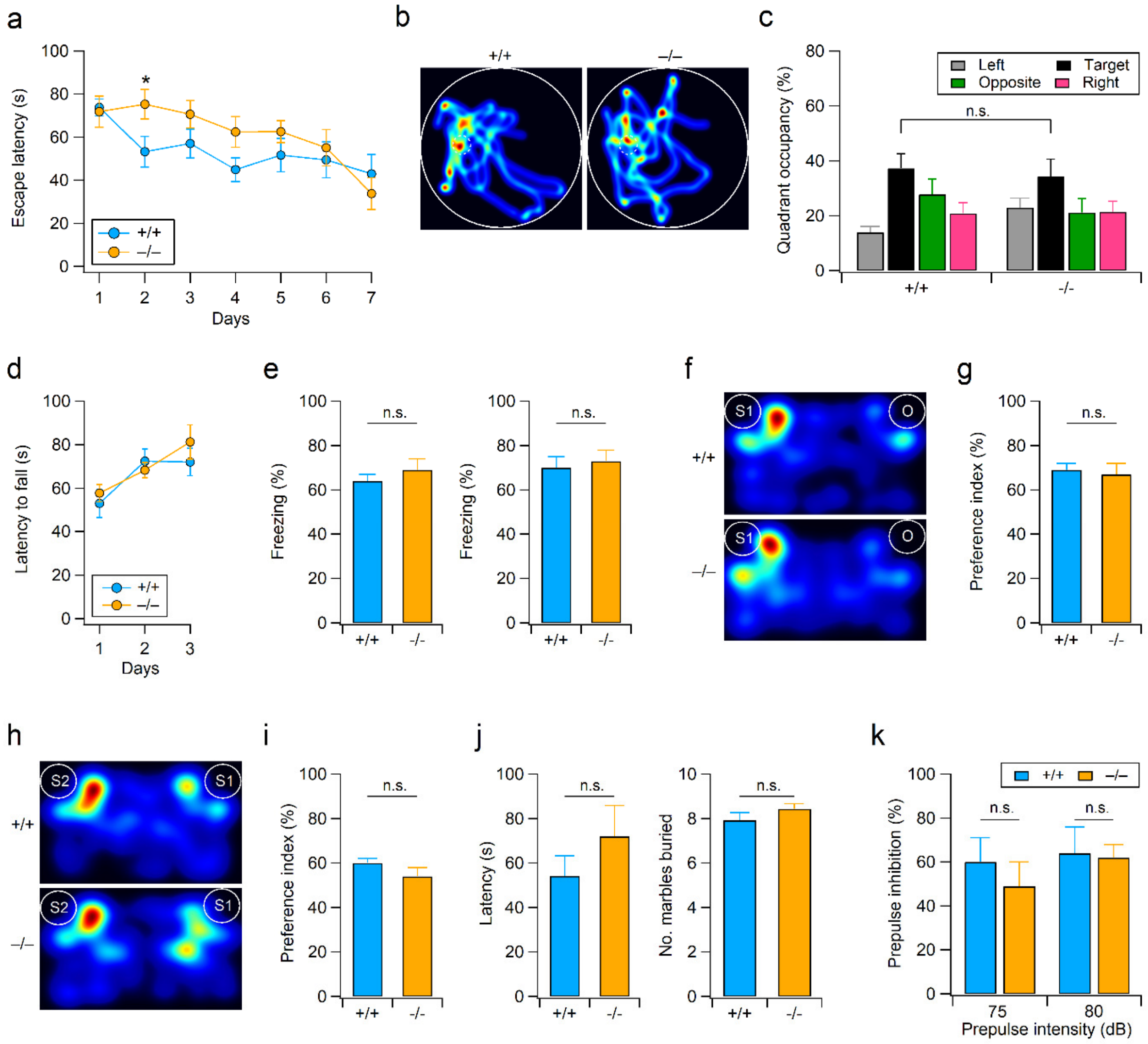


**Supplementary Fig. 4: Normal learning, social interaction and sensory gating in RalBP1$^{-/-}$ mice.** (a-e) Intact hippocampus-dependent learning in RalBP1$^{-/-}$ mice. (a) The escape latencies were averaged from 4 trials on each day during the Morris water maze training. *P < 0.05 by Student t-test. (b) Representative swim paths of mice during the probe trials (60 s) after completion of 7 days of training. (c) The percentage of time spent in each quadrant during the probe trials. (d) RalBP1$^{-/-}$ mice display normal motor coordination and motor learning. The latencies to falling from the accelerating rotarod are plotted against the training days. (e) Quantification of context-specific (left) and cue-specific (right) freezing behaviors measured 24 h after the fear conditioning training. (f-i) Normal social approach and social novelty recognition in RalBP1$^{-/-}$ mice. (f) Representative activity traces of WT (top) and RalBP1$^{-/-}$ (bottom) mice during the three-chamber social approach test. (g) Both genotypes spent more time in the social chamber containing a stranger mouse (S1) than the chamber containing an inanimate object (O). Preference index (%) = Time sniffing the stranger / (Time spent sniffing the stranger + Time spent sniffing the object) × 100. (h) Representative heat maps show the location of WT (top) and RalBP1$^{-/-}$ (bottom) mice during the social novelty recognition test. (i) Quantification of preference for a new stranger mouse (S2). Preference index (%) = Time spent sniffing the new stranger (S2) / (Time spent sniffing the new stranger (S2) + Time spent sniffing the familiar (S1)) × 100. (j) Latency (left) to start marble burying and the number (right) of marble buried by WT and RalBP1$^{-/-}$ mice are summarized. (k) Prepulse inhibition (PPI) of acoustic startle response at two different prepulse intensities (75 and 80 dB).

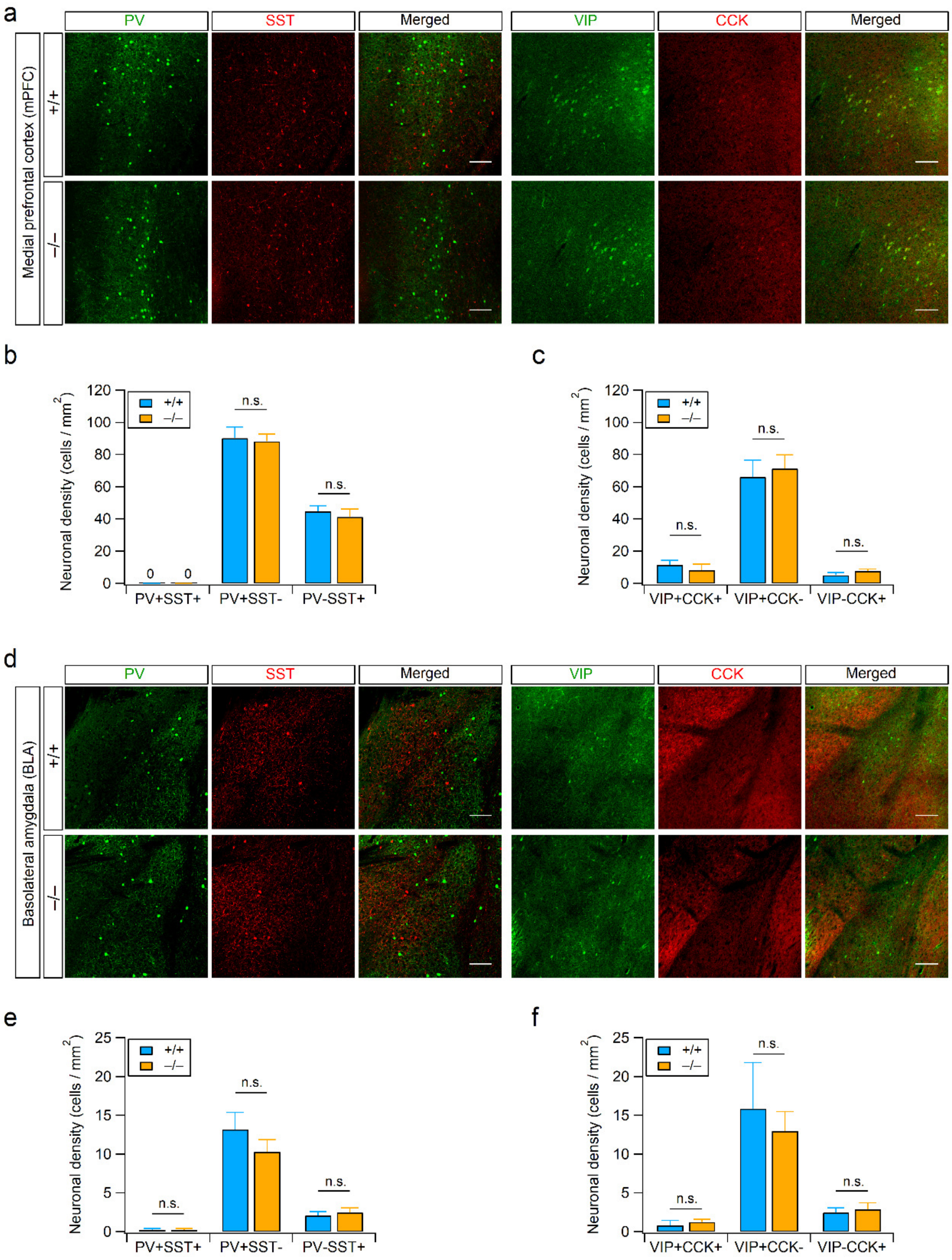


**Supplementary Fig. 5: Normal density of interneurons in the RalBP1$^{-/-}$ medial prefrontal cortex (mPFC) and basolateral amygdala (BLA).** (a) Immunohistochemical staining for parvalbumin (PV), somatostatin (SST), vasoactive intestinal peptide (VIP) and cholecystokinin (CCK) in the mPFC from WT and RalBP1$^{-/-}$ mice. (b) RalBP1$^{-/-}$ mice exhibit normal population of PV and SST neurons in the mPFC. (c) Quantification of VIP- and/or CCK-expressing neurons in the mPFC. (d) The distributions of PV, SST, VIP and CCK were examined by immunohistochemistry in the BLA from WT and RalBP1$^{-/-}$ mice. (e) Neurons expressing PV and/or SST in the BLA are unaltered by RalBP1 mutation. (f) Quantification of VIP- and/or CCK-positive neurons in the BLA. (a and d) Scale bars, 100 µm.

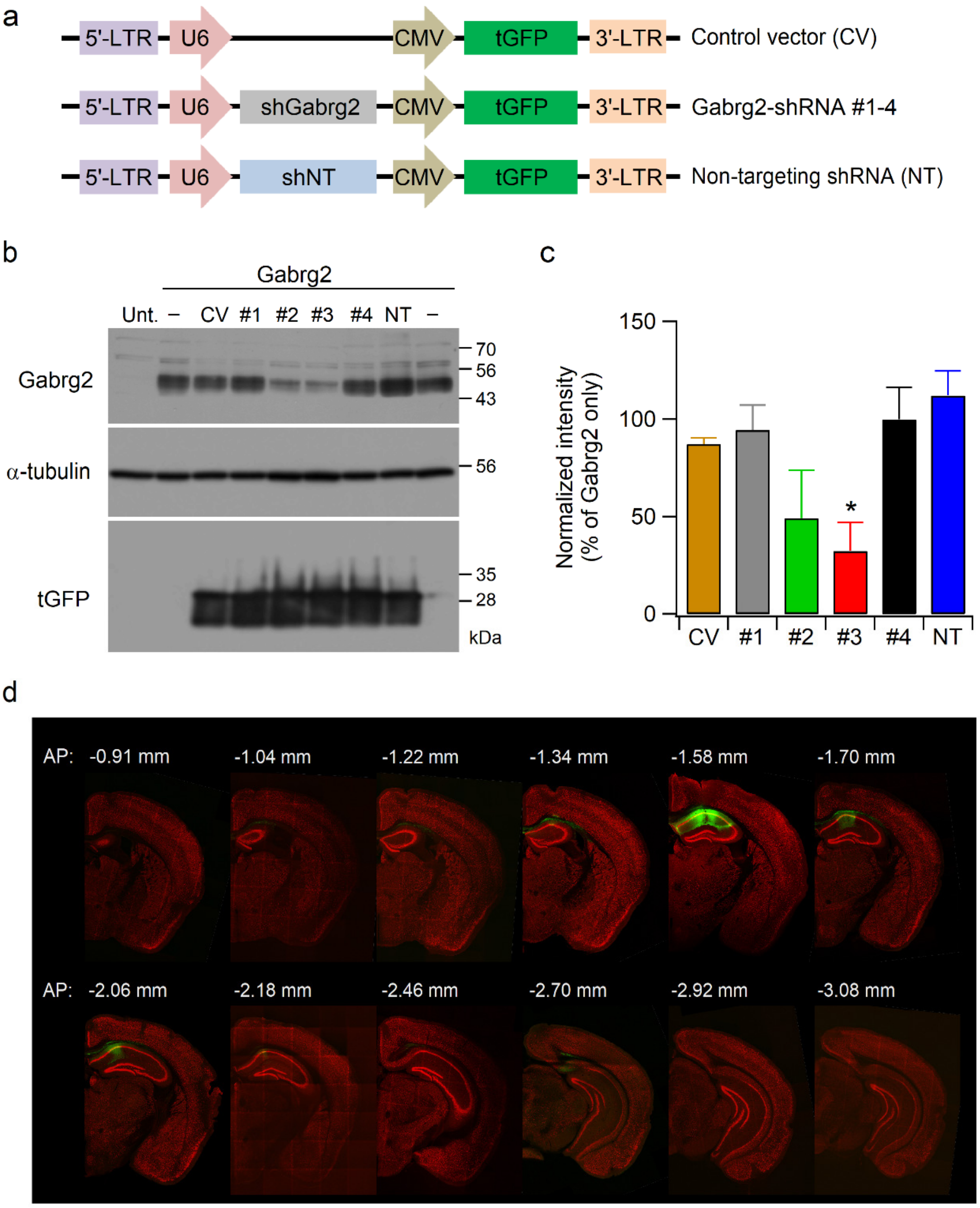


**Supplementary Fig. 6: Efficiencies of Gabrg2 knockdown in HEK293T cells and lentivirus infection in the CA1 area of the mouse brain.** (a-c) Western blot analysis of Gabrg2 knockdown efficiencies of four different shRNA sequences (sh#1-4) in HEK293T cells. (a) Schematic representation of the shRNA constructs. (b) Representative western blots for Gabrg2, α-tubulin and TurboGFP (tGFP) obtained from HET293T cell lysates at 72 h post-transfection. Control vector (CV) expresses tGFP without a hairpin sequence. The non-targeting (NT) shRNA contains a tGFP expression cassette and the hairpin with surrounding sequences that do not overlap with any known mouse gene (see Methods for details). α-tubulin and tGFP were used as loading and transfection controls, respectively. Unt., untransfected control HEK293T cells. (c) Quantification of the Gabrg2 knockdown efficiency in transfected cell lines. (d) Series of immunofluorescent images of coronal sections co-stained with eGFP (green) and NeuN (red) show the distribution of lentivirus-infected cells (green) in the mouse brain. Numbers indicate anterior/posterior (AP) coordinates to Bregma.

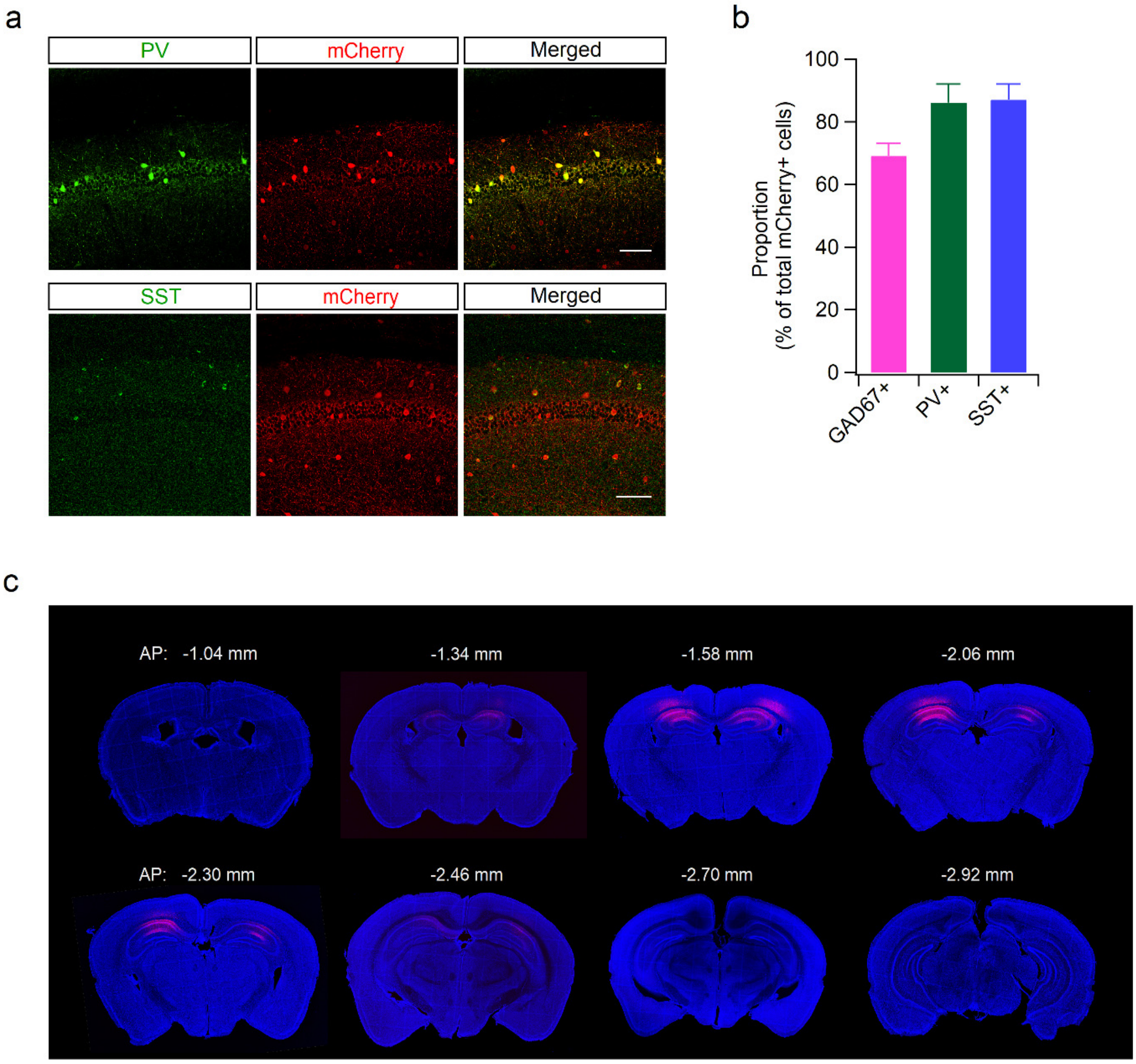


**Supplementary Fig. 7: Expression of mCherry in the PV- and SST-expressing neurons in the hippocampal CA1 area.** (a) Hippocampal section from Vgat-Cre mice infected with DiO-DREADDs-mCherry was double immunostained with PV/mCherry (top) or SST/mCherry (bottom) antibodies. Scale bars, 100 µm. (b) Quantification of mCherry expressing cells in the hippocampal CA1 regions. mCherry signals were detected in 69%, 86%, and 87% of GAD67-, PV-, and SST-positive cells, respectively. N = 16 slices from 5 mice (GAD67), 7 slices from 3 mice (PV), and 6 slices from 3 mice (SST). (c) Series of mCherry immunofluorescent images of coronal brain sections show the distribution of cells expressing mCherry (red) in the Vgat-Cre mouse. mCherry signals are mainly located in the CA1 subfields of the dorsal hippocampus. DAPI (blue) was used for the identification of brain regions. Numbers indicate anterior/posterior (AP) coordinates to Bregma.

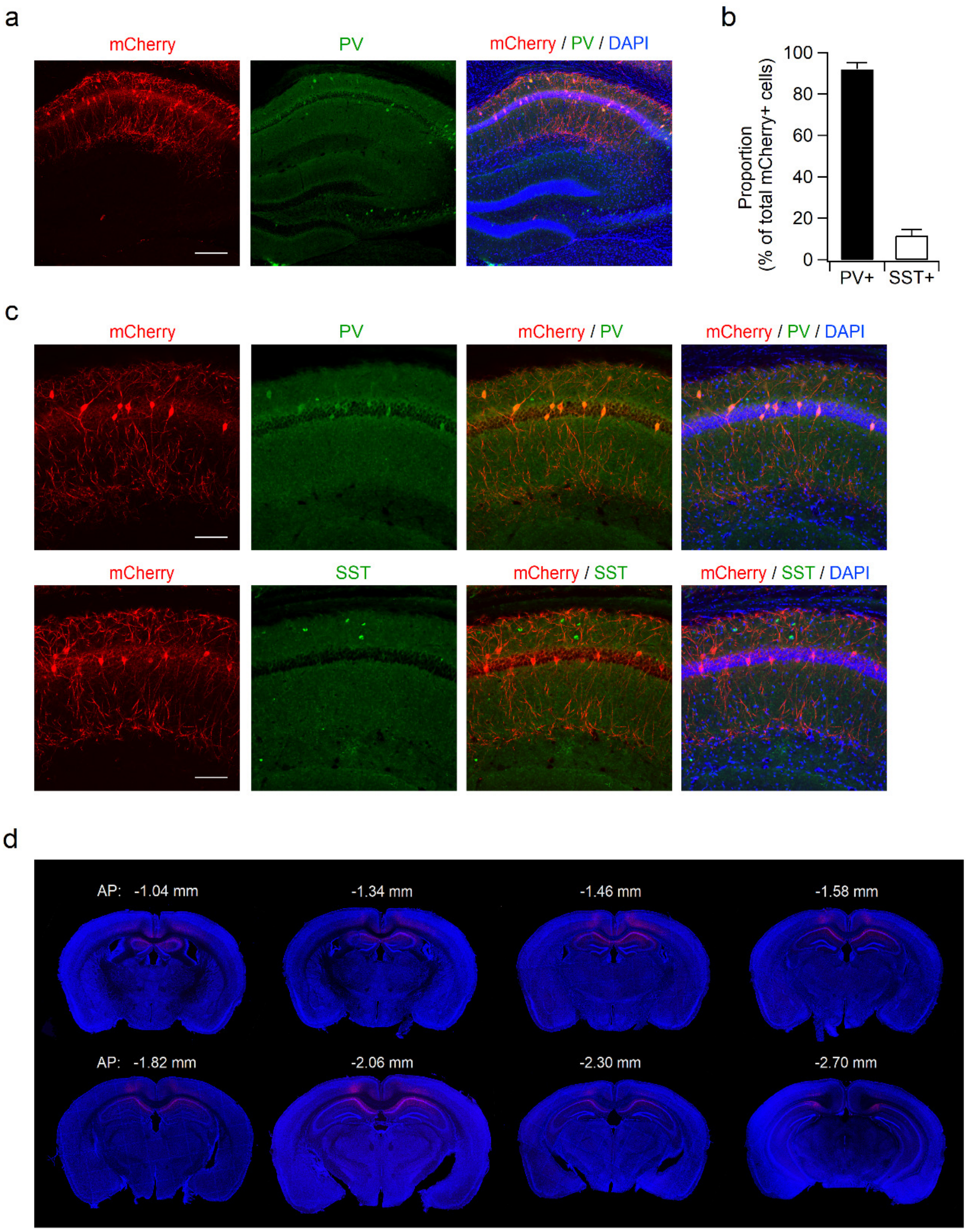


**Supplementary Fig. 8: mCherry was mainly localized in the PV-positive cells in the hippocampus of PV-Cre mice.** (a) Sample images showing the distribution of mCherry (red) and PV (green) in the dorsal hippocampus of PV-Cre mice infected with DiO-DREADDs-mCherry. Scale bar, 200 μm. (b) mCherry signals in the CA1 area were detected in 92% and 12% of PV- and SST-positive cells, respectively. N = 8 slices from 4 mice. Scale bars, 100 μm. (c) Co-immunostainings of hippocampal sections with PV/mCherry (top) and SST/mCherry (bottom) show preferential expression of mCherry in the PV-expressing cells. (d) The series of mCherry immunofluorescent images of coronal brain sections showing the distribution of mCherry (red) in the PV-Cre mouse. Numbers indicate anterior/posterior (AP) coordinates to Bregma. DAPI (blue) was used for identification of brain regions and structures (a,c,d).

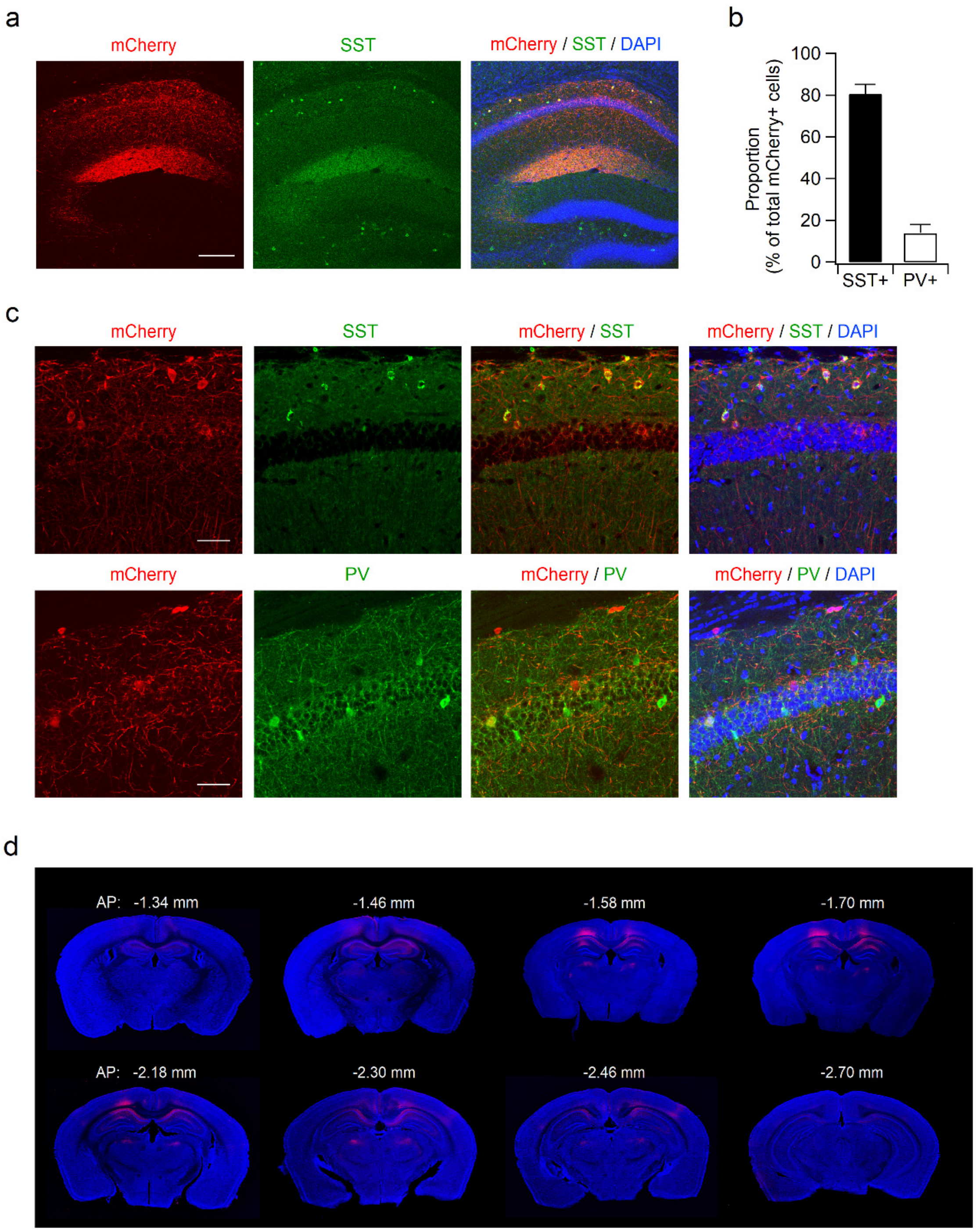


**Supplementary Fig. 9: mCherry signals are mainly detected in the SST-immunoreactive cells by stereotaxic injection of AAV-DiO-DREADDs-mCherry/AAV-SST-Cre mixture.** (a) Sample images showing the distribution of mCherry (red) and SST (green) in the dorsal hippocampus of WT mice infected with a mix of AAV-DiO-DREADDS-mCherry and AAV-SST-Cre. Scale bar, 200 μm (b) Quantification of mCherry-expressing cells shows the predominant expression of mCherry in SST-positive cells (81%) than PV-positive cells (13%) in the hippocampal CA1 area. N = 6 slices from 3 mice for each cell type. (c) Co-immunostainings of hippocampal sections with SST/mCherry (top) and PV/mCherry (bottom) reveal predominant expression of mCherry in SST-expressing cells. Scale bars, 50 μm. (d) The distribution of mCherry-expressing cells in the mouse brain infected with a mix of AAV-DiO-DREADDs-mCherry and AAV-SST-Cre was determined by immunohistochemical staining with mCherry antibodies. DAPI (blue) was used to identify brain regions and structures (a, c, d).

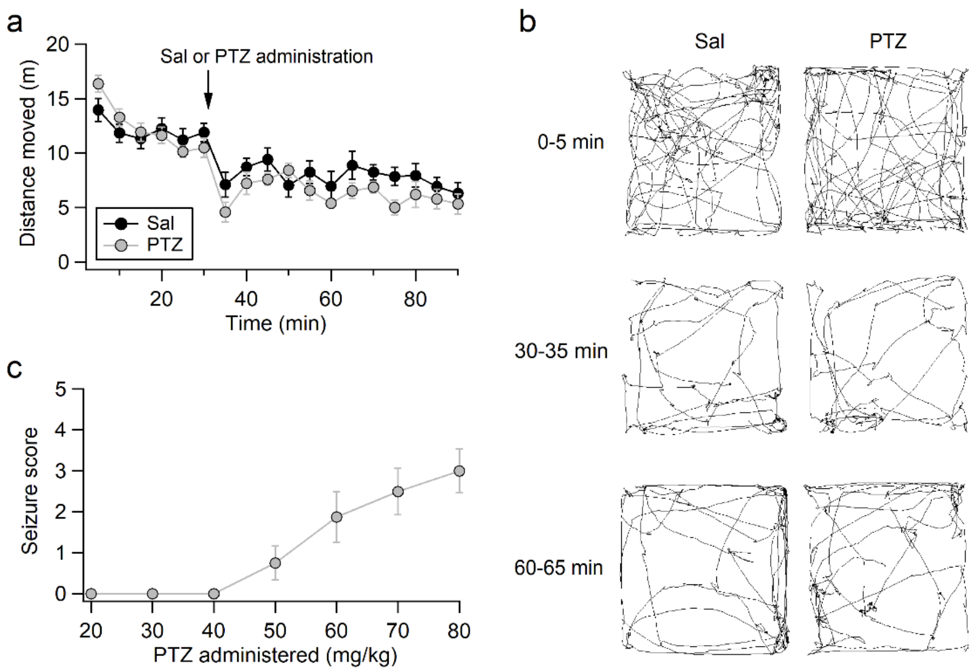


**Supplementary Fig. 10: Low-dose administration of pentylenetetrazol (PTZ) does not induce seizure activity or hyperlocomotion.** (a) Mice received PTZ (20 mg/kg) or saline (Sal) 30 min after baseline activity recording during the 90-min open-field test. (b) Sample path recordings during the first 5 min of the open field test, and immediately after (30-35 min) and 30 min after (60-65 min) drug administration. (c) Dose-response curves of the effect of different PTZ doses on behavioral seizures in male mice. Animals were monitored for 1 h after the intraperitoneal injection of PTZ. The severity of the behavioral seizures was scored as follows: score 0, no seizure; score 1, head nodding; score 2, clonic jerks, sporadic shaking; score 3, full-body spasms, Straub tail, rearing; score 4, shrieking, jumping, falling; score 5, violent convulsions, death. N = 8-10 mice for each dose.

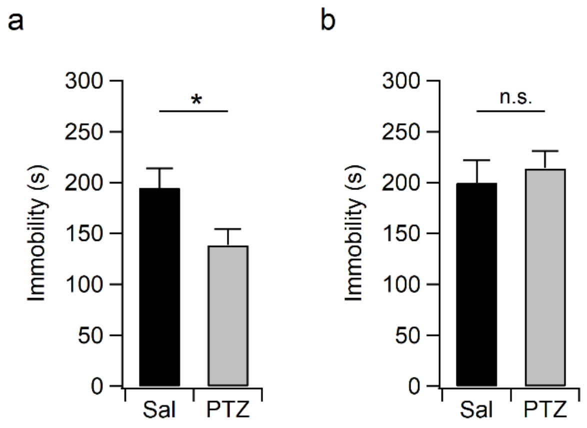


**Supplementary Fig. 11: Single PTZ administration induces rapid but not long-lasting antidepressant behaviors**. (a, b) Bar graphs show total immobility time during the 6-min-TST measured 30 min (a) and 48 h (b) after PTZ injection (20 mg/kg, i.p.).

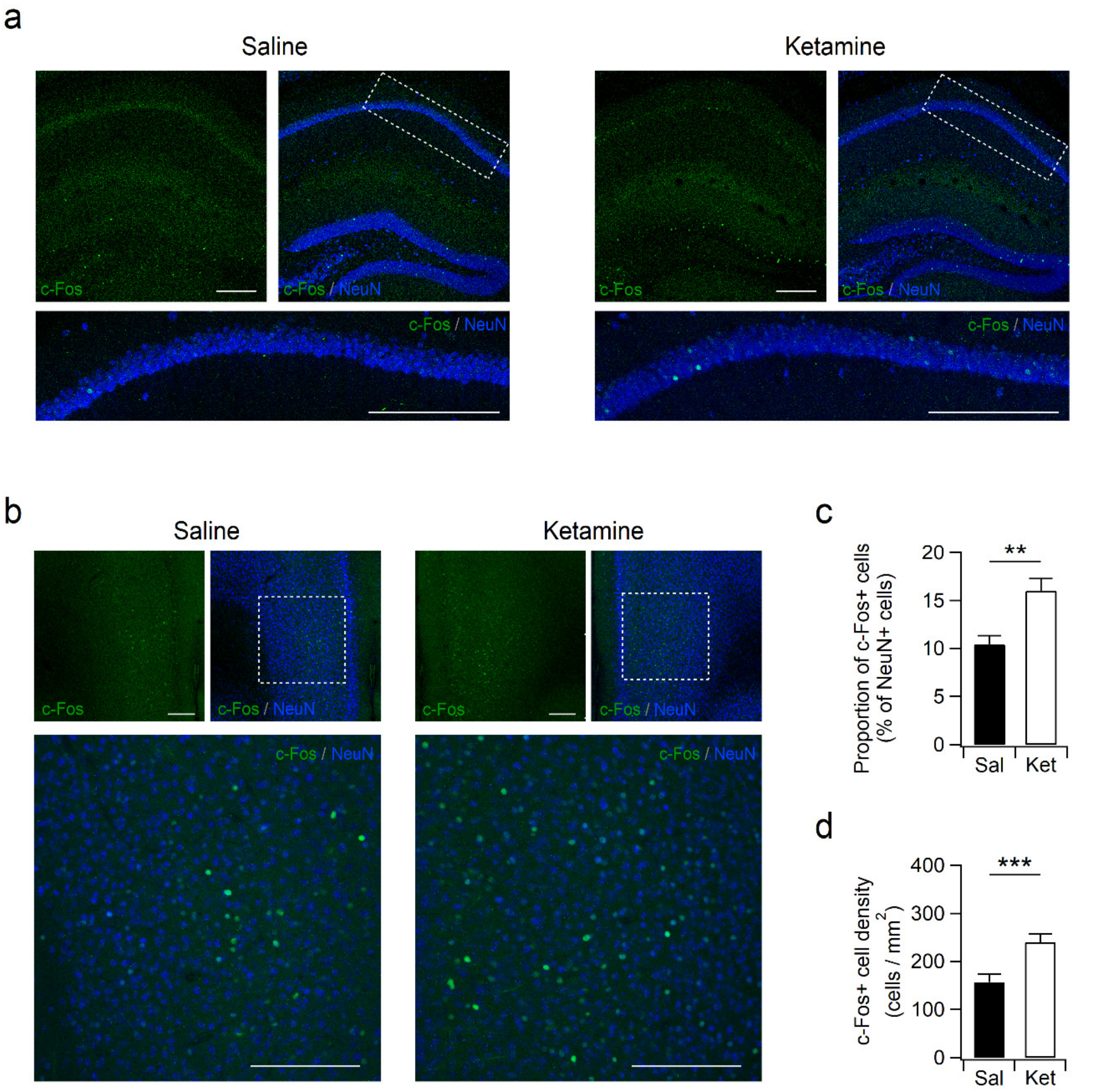


**Supplementary Fig. 12: Enhanced c-Fos expression in the hippocampus and mPFC of ketamine-treated mice.** (a) Coronal hippocampal sections from saline- or ketamine-treated mice were co-immunostained with c-Fos (green) and NeuN (blue) antibodies. Bottom images show the white dotted box regions magnified from top images. Sections were prepared 1 h after saline or ketamine injection. (b) Ketamine induced rapid enhancement in c-Fos (green) expression in the mPFC. NeuN (blue) was used to identify neurons in the mPFC. Bottom shows magnified images indicated by the dotted white boxes in the upper panels. Scale bars, 200 μm (a,b). (c) The number of cells expressing c-Fos were normalized to the total number of NeuN+ cells in the mPFC. (d) Bar graphs represent the density of cells expressing c-Fos in the mPFC of mice receiving saline or ketamine. N = 6 slices from 3 mice (c,d).

Fig. 2d

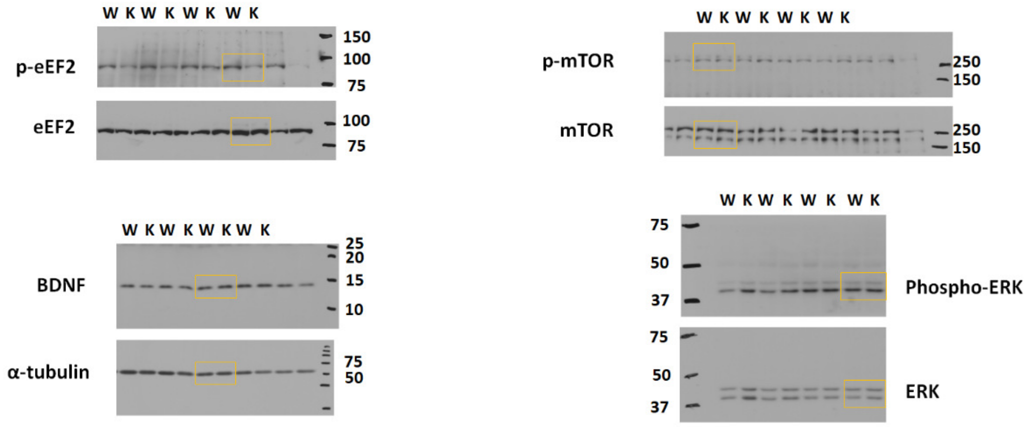


Fig. 6d

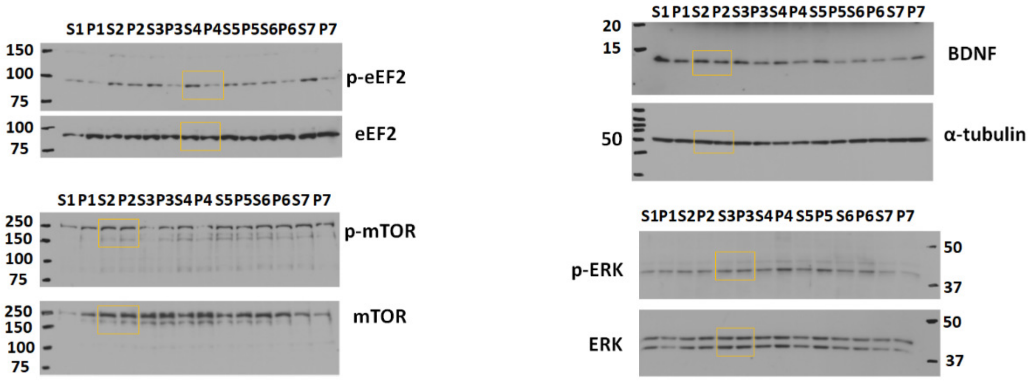


Fig. 6h

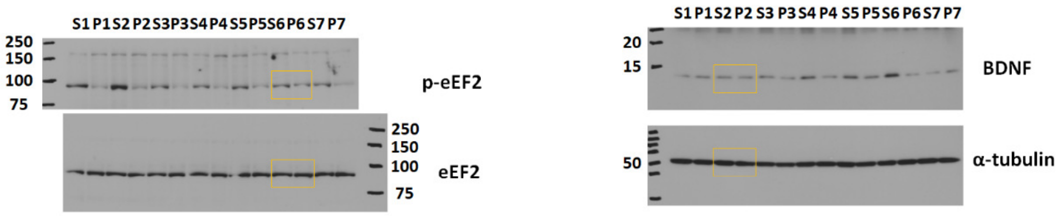


Fig. 7h

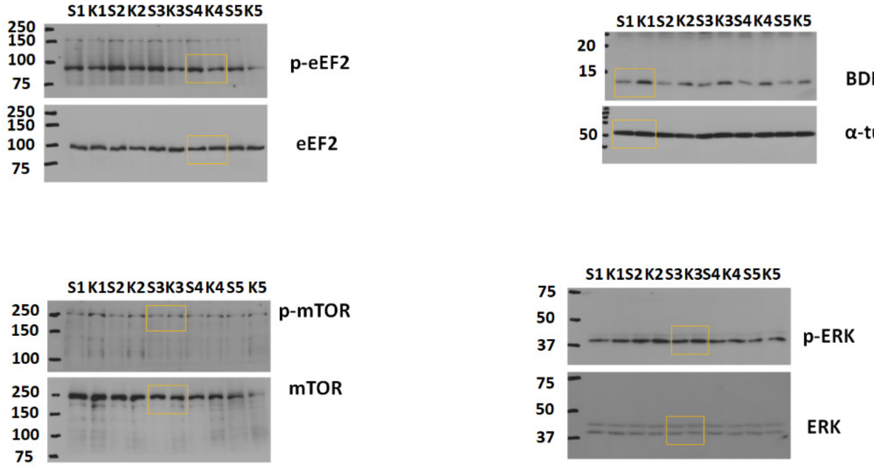


**Supplementary Fig. 13: Uncropped western blot images presented in the main figures.**
